# The effects of seasonal anoxia on the microbial community structure in demosponges in a marine lake (Lough Hyne, Ireland)

**DOI:** 10.1101/2020.09.09.290791

**Authors:** Astrid Schuster, Brian William Strehlow, Lisa Eckford-Soper, Rob McAllen, Donald Eugene Canfield

## Abstract

Climate change is expanding marine oxygen minimum zones (OMZs), while anthropogenic nutrient input depletes oxygen concentrations locally. The effects of deoxygenation on animals are generally detrimental; however, some sponges (Porifera) exhibit hypoxic and anoxic tolerance through currently unknown mechanisms. Sponges harbor highly specific microbiomes, which can include microbes with anaerobic capabilities. Sponge-microbe symbioses must also have persisted through multiple anoxic/hypoxic periods throughout Earth history. Since sponges lack key components of the hypoxia-inducible factor (HIF) pathway responsible for hypoxic responses in other animals, it was hypothesized that sponge tolerance to deoxygenation may be facilitated by its microbiome. To test this hypothesis, we determined the microbial composition of sponge species tolerating seasonal anoxia and hypoxia *in situ* in a semi-enclosed marine lake, using 16S rRNA amplicon sequencing. We discovered a high degree of cryptic diversity among sponge species tolerating seasonal deoxygenation, including at least nine encrusting species of the orders Axinellida and Poecilosclerida. Despite significant changes in microbial community structure in the water, sponge microbiomes were species specific and remarkably stable under varied oxygen conditions, though some symbiont sharing occurred under anoxia. At least three symbiont combinations, all including large populations of *Thaumarchaeota*, corresponded with deoxygenation tolerance, and some combinations were shared between distantly related hosts. We propose hypothetical host-symbiont interactions following deoxygenation that could confer deoxygenation tolerance.

**Importance:** The oceans have an uncertain future due to anthropogenic stressors and an uncertain past that is becoming clearer with advances in biogeochemistry. Both past and future oceans were, or will be, deoxygenated compared to present conditions. Studying how sponges and their associated microbes tolerate deoxygenation provides insights into future marine ecosystems. Moreover, sponges form the earliest branch of the animal evolutionary tree and they likely resemble some of the first animals. We determined the effects of variable environmental oxygen concentrations on the microbial communities of several demosponge species during seasonal anoxia in the field. Our results indicate that anoxic tolerance in some sponges may depend on their symbionts, but anoxic tolerance was not universal in sponges. Therefore, some sponge species could likely outcompete benthic organisms like corals in future, reduced-oxygen ecosystems. Our results support the molecular evidence that sponges and other animals have a Neoproterozoic origin, and that animal evolution was not limited by low-oxygen conditions.

## Introduction

Ocean anoxia and hypoxia have become major stressors for many marine organisms. Indeed, oxygen-minimum zones (OMZs) and coastal hypoxic areas will most likely expand in the future (1–5), leading to habitat and biodiversity losses (5–8), where oxygen depletion is caused by both natural and anthropogenic influences (9, 10). While motile species can escape such hypoxic/anoxic areas, many sessile organisms like sponges (Porifera) and corals (Cnidaria) must either cope with these extremes or suffer mass mortalities as observed in Tropical dead zones (11). Nevertheless, the lethal thresholds of, and potential adaptations to, deoxygenation are understudied in these organisms (12), particularly in sponges.

Sponges are common, cosmopolitan filter feeders that pump water through their bodies to filter and ingest nutrients and microorganisms (13, 14). Previous *ex situ* experiments have shown that sponges, including *Geodia barretti* (15), *Tethya wilhelma* (16), *Halichondria panicea* (17), *Haliclona pigmentifera* (18) and *Vazella pourtalesii* (19), have a high tolerance of hypoxia. For instance, *T. wilhelma* can maintain normal transcription at ∼0.5 µM O_2_ (16), despite lacking key components of the hypoxia-inducible factor (HIF) pathway, which regulates hypoxic responses in other invertebrates (16). However, *T. wilhelma, H. panacea*, and *H. pigmentifera* became stressed, sometimes fatally, during anoxia (16–18). Some marine sponge species also tolerate hypoxia and even anoxia in their natural environment (20–24), and gemmules from freshwater sponges can survive months of anoxia (25). Nevertheless, hypoxia-induced mortality was reported *in situ* for the demosponges *Aaptos simplex* and *Homaxinella amphispicula* (21), so responses are likely species-specific. The current global distribution and depth range of sponges (26) overlaps with that of hypoxic areas worldwide (2), indicating that many sponges may even thrive in hypoxic environments, perhaps due to limited competition. Therefore, some sponges have likely developed alternative adaptation strategies to tolerate variable and low-oxygen conditions independent of the HIF pathway.

Sponges can harbor stable and sometimes diverse microbial communities that can constitute up to 50% of their biomass (27, 28); together, the host and microbiome are referred to as the sponge ‘holobiont’ (29). Sponge holobionts play a key role in marine ecosystems, contributing to reef formation, benthic-pelagic nutrient coupling, and biogeochemical cycling (30–32). Some of these sponge-microbe associations are highly stable under environmental stressors, including elevated temperature (33–35), eutrophication and sedimentation (36), and some symbionts are sponge-specific, meaning they occur negligibly in the environment (37). Stable sponge-microbe associations were also found across large geographic distances (38). However, microbial communities across sponge species substantially vary in diversity, structure and abundance (39), and there can be selection for divergent microbiomes, even among related sponge lineages (40).

Within a specific holobiont, symbiotic microbes may increase sponge fitness by providing food (through carbon fixation or direct ingestion of symbionts), recycling of nutrients and waste products, and/or the production of secondary metabolites for predator defense or other functions (30, 31). Many sponge species form symbioses with *Nitrosopumilus*-like ammonia-oxidizing *Archaea* (AOA) and/or *Nitrospira* sp. nitrite-oxidizing *Bacteria* (NOB) (19, 40–44). Each of these prokaryotic groups utilize sponge waste ammonia for nitrification. Similar microbes are active in hypoxic waters and contribute to biogeochemical cycling processes (45). For example, *Nitrospira* sp. accounted for 9% of the total microbial community in the OMZ of the Benguela upwelling system (46). Also, *Nitrosopumilacea* sp., a widespread and dominant AOA in many OMZs, plays a significant role in ammonium oxidation therein (47, 48). Thus, many of these common sponge symbionts could be adapted to hypoxia, and it was suggested that the presence of AOA symbionts, and other facultative anaerobes, within a glass sponge signified holobiont adaptation to mild, persistent environmental hypoxia (19). Furthermore, some symbionts have anaerobic metabolisms, including sulfate reducers (49), denitrifiers, and anaerobic ammonium oxidizers (anammox bacteria) (50). Localised and widespread anoxia is common within sponge tissue, even within oxygenated environments, due to pumping cessation, and this allows for anaerobic metabolism of the symbionts (49). Such anaerobic metabolism may be crucial for nutrient cycling.

The low-oxygen tolerance of the sponge holobiont was probably crucial throughout evolutionary history. The symbiosis of sponges with microbes dates back hundreds of millions of years (51–53), indicating adaptations of holobionts to environmental changes and strong coevolution throughout Earth’s history, including long periods of anoxia/hypoxia (54–58). For example, hypercalcified sponges likely survived the late and end-Permian mass extinction and deoxygenation because of their oxygenic cyanobacterial symbionts (59). Based on morphology (60) and phylogeny (61), we assume that the last common ancestor of metazoans was sponge-like. Therefore, the low-oxygen tolerance in early metazoans, similar to that of modern sponges, could have permitted their evolution under lower-oxygen concentrations (1–20% of present levels, (62)) in the Neoproterozoic Era (16, 63, 64). Although the symbiont status of early metazoans is unknown, sponges evolved in a world rich in microbes (65), and hypoxic tolerance of modern sponge holobionts could date back to these associations.

Based on the microaerophilic and anaerobic metabolisms of some sponge symbionts (66, 67), and the evolutionary importance of hypoxic tolerance, we hypothesized that specific microbes or microbial compositions help sponges survive in low-oxygen or even anoxic environments. To test this hypothesis, sponge, sediment and water samples from a semi-enclosed marine lake (Lough Hyne, Ireland, Fig. 1A) were sampled and analyzed using 16S amplicon sequencing to determine microbial community structure and molecular barcoding to identify sponge species. Lough Hyne is characterized by its unique ecological, physical and chemical conditions (see 68). This lough was a freshwater basin until around 4,000 years ago (69), but became a marine basin due to rising sea levels and intrusion from the Atlantic Ocean over a shallow sill. Water retention in the lough is between 14–41 days (70, 71). In the summer, a seasonal anoxia/hypoxia layer forms at a depth of approximately 25 m. Water masses beneath this layer become anoxic and enriched in H_2_S, NH_4_^+^ and Mn^2+^ (68, 69). Lough Hyne also harbors a remarkably high biodiversity of sponges, and some of these species were reported to survive seasonal anoxia (72), making it a natural laboratory to study the effects of variable oxygen on sponges-microbiome associations.

**Fig. 1.**
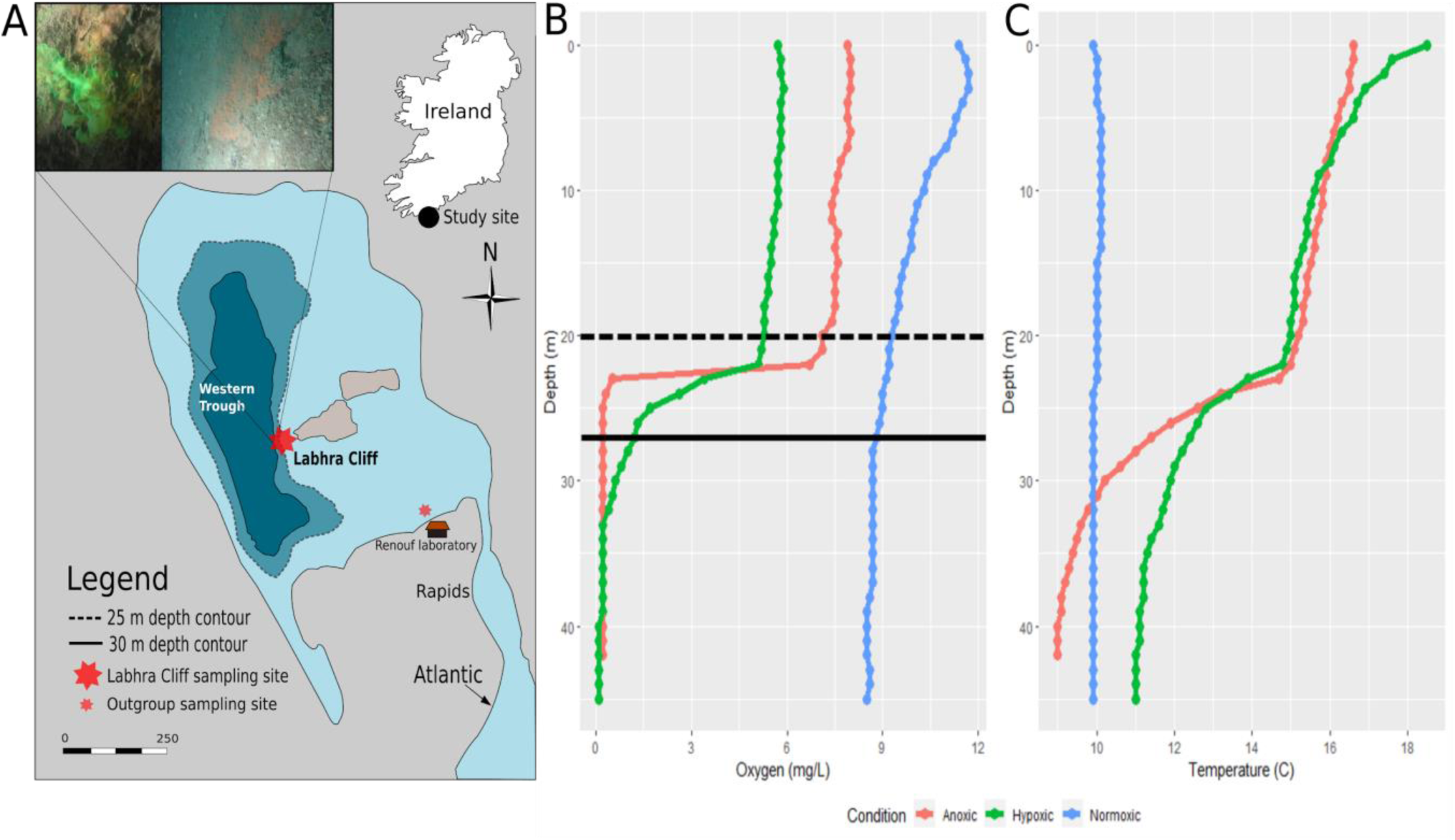
Map of Lough Hyne with sampling sites, oxygen and temperature profiles taken from the middle of the Western Trough in Lough Hyne. A. Map of Lough Hyne showing sampling sites and *in situ* pictures of encrusting sponges at 27 m depth. The picture upper left shows the sponge actively pumping as visualized using fluorescein dye under hypoxic conditions and the right picture shows a sponge without dye. B. Dissolved oxygen concentrations versus depth for the three different sampling trips, corresponding to three different oxygen conditions at ∼27 m (solid line): July 2018, anoxic (red), July 2019, hypoxic (green) and April 2019, normoxic (blue). Samples were collected at ∼27 m (solid line, below the thermocline) and at ∼20m (dashed line, above the thermocline) for comparison across time and oxygen condition. C. Temperature versus depth during the different sampling oxygen conditions.

Samples were taken above and below the thermocline, and during different seasons, to allow a comparison of a wide range of oxygen conditions from normoxia to anoxia. In addition, three sponge species were sampled outside the seasonally anoxic site for comparison (Fig. 1A). This study aimed to answer the following questions: (1) what sponge species survive seasonal anoxia/hypoxia in Lough Hyne, (2) is there a symbiont composition that corresponds to deoxygenation tolerance and finally, (3) what implications does sponge holobiont deoxygenation tolerance have for early animal evolution in low-oxygen environments and in future oceans? To our knowledge, this is the first study to comparatively investigate the microbial community composition of sponges under different *in situ* oxygen levels.

## Results

### Physical data

Sampling was performed in July 2018, April 2019 and July 2019 when oxygen conditions at Labhra Cliff (Fig. 1A), between 25–30 m deep, were respectively anoxic, normoxic and hypoxic (Fig. 1B). The hypoxic conditions in July 2019 were anomalous compared to the normally anoxic summer conditions below 25 m at this site and likely occurred due to heavy storm and wind mixing of waters in the lough. These storm events increased mixing across the thermocline and pushed resultant anoxia to a depth of at least 33 m, which is below the depth of the cliff/dive site (Fig. 1B, C). Measurements taken by CTD in the center of the Western Trough (4 casts to 40 m) and at Labhra Cliff (2 casts to ∼30 m) indicated that total anoxia occurred at 40 m depth (Fig. S1B), while conditions between 25–30 m were still ‘hypoxic’ (50–150 uM, Fig. S1A). Although CTD measurements were not performed in July 2018, the total depletion of oxygen in the ‘anoxic’ condition was verified by the presence of sulfide in the water which cannot persist in the presence of oxygen. Sulfide concentrations in two samples were 0.90 µM and 0.19 μM. When these samples were quickly frozen, sulfide was not stabilized as ZnS acetate until they were returned to the lab, so the actual sulfide concentrations were likely higher. Furthermore, following anoxic collections, all water samples and equipment including SCUBA gear had the characteristic smell of sulfide. The presence of photosynthetic pigments (Fig. S2) indicated that there was a potential food source for sponges regardless of oxygen condition.

### Diversity of sponges surviving seasonal anoxia/hypoxia

Using molecular barcoding, 30 demosponge specimens were identified from the Labhra cliff site. The majority of specimens belonged to the Family Raspailiidae (Subfamily Raspailiinae) of the Order Axinellida except for *Mycale* sp., a non-carnivorous poecilosclerid sponge collected under anoxic conditions (Fig. 2 and Fig. S3), but all were visually similar (orange-red, encrusting) *in situ*. Within the Axinellida, species of the genera *Eurypon, Endectyon, Raspaciona* and *Hymeraphia* (Fig. 2) were sampled. This includes seven species that are definitely exposed to seasonal anoxia/hypoxia, namely *Eurypon* sp.2 (n=14), *Eurypon clavigerum* (n=2), *Eurypon* cf. *cinctum* (n=1), *Hymeraphia stellifera* (n=8), *Endectyon* sp.1 (n=2), *Endectyon* sp.2 (n=2) and *Raspaciona* sp. (n=1) (Fig. 2 and Fig. S4).

**Fig. 2.**
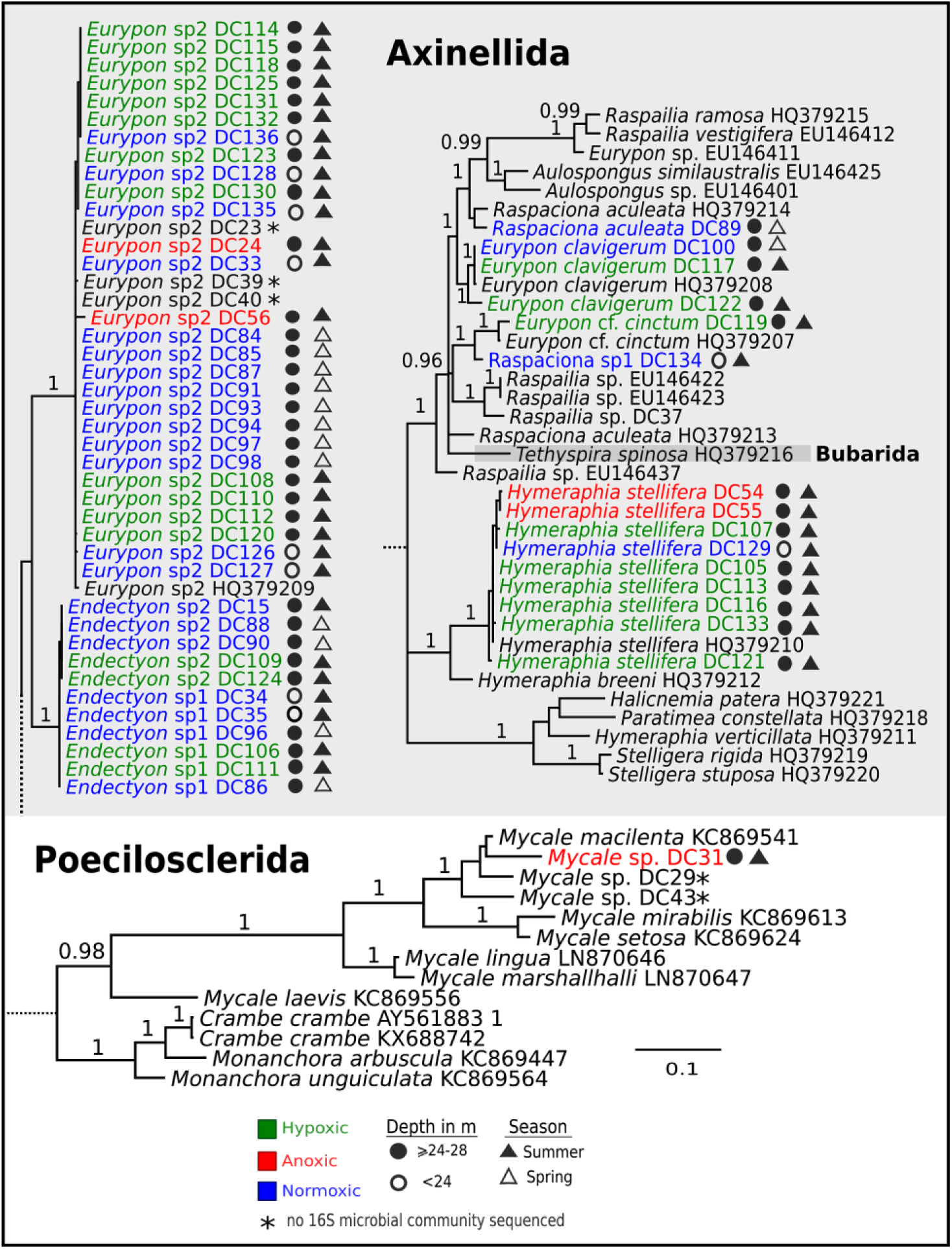
28S Bayesian Inference (BI) phylogeny of sponges from Lough Hyne (indicated by DC numbers after taxa names). For visualization, subtrees were pruned from the complete phylogenetic tree (see Fig. S3). Posterior Probability values are given above branches for >0.95. See Fig. S4 for *cox1* BI phylogeny.

However, given the challenging sampling conditions at this site in the lough (see M&M), it is possible that other species such as *Amphilectus* spp., *Rhizaxinella* sp. and *Hymeniacidon perlevis*, collected above 24 m, may also be present in the anoxic/hypoxic layers but were simply not sampled during our dives. Notably, all species sampled from below 24 m were encrusting in their growth form.

Systematically, the genera *Eurypon, Hymeraphia* and *Endectyon* were polyphyletic (Fig. 2 and Fig. S3). Both markers (28S and *cox1*) indicated that *Eurypon* spp. most likely represents a complex of species, which will be taxonomically resolved in the future using more informative markers and holotypes. All sponges exposed to fluorescein dye at 27 m were pumping, under both normoxic and hypoxic conditions (Fig. 1A). This assay was not conducted under anoxia due to logistical constraints.

### Microbial communities

A total of 89 sponge, water and sediment samples were collected from Labhra cliff in which 4677 OTUs were identified. An additional 114 OTUs were present in the 14 outgroup samples, consisting of sponges sampled in normoxic waters away from Labhra Cliff (Fig. 1). For all samples, the average Shannon index was 3.1 and approximately 500 OTUs were identified in each sample. The data was filtered into a ‘focused subset’ to only include species with replication across all oxygen conditions. The focused subset included the sponges *Eurypon* sp.2 and *Hymeraphia stellifera* as well as water and sediment samples, but notably, no sediment sample under anoxic conditions were collected. In total, this dataset comprised 55 samples, containing 4518 OTUs (∼600 per sample) with an average Shannon index of ∼3 per sample. A PERMANOVA performed on the focused subset indicated that although there were some microbial community shifts based on oxygen condition and the interaction of oxygen condition and sample type, these factors only accounted for 7.33 and 7.84% of the variance in OTUs within the dataset, respectively (Table 1). Most of the variance (67.8%, Table 1) in microbial community structure came from the type of sample, i.e. *E*. sp.2, *H. stillefera*, water or sediment, which was also clear from a principal component analysis (PCA, Fig. 3A).

**Table 1.** PERMANOVA results for the focused subset after 1000 permutations.

| Factor | df | F | R<sup>2</sup> | p |
| --- | --- | --- | --- | --- |
| Oxygen | 2 | 10.2 | 0.0777 | <0.001 |
| Sample type | 3 | 59.6 | 0.678 | <0.001 |
| Oxygen x sample type | 5 | 4.03 | 0.0765 | <0.001 |
| Residuals | 44 |  |  |  |

**Fig. 3.**
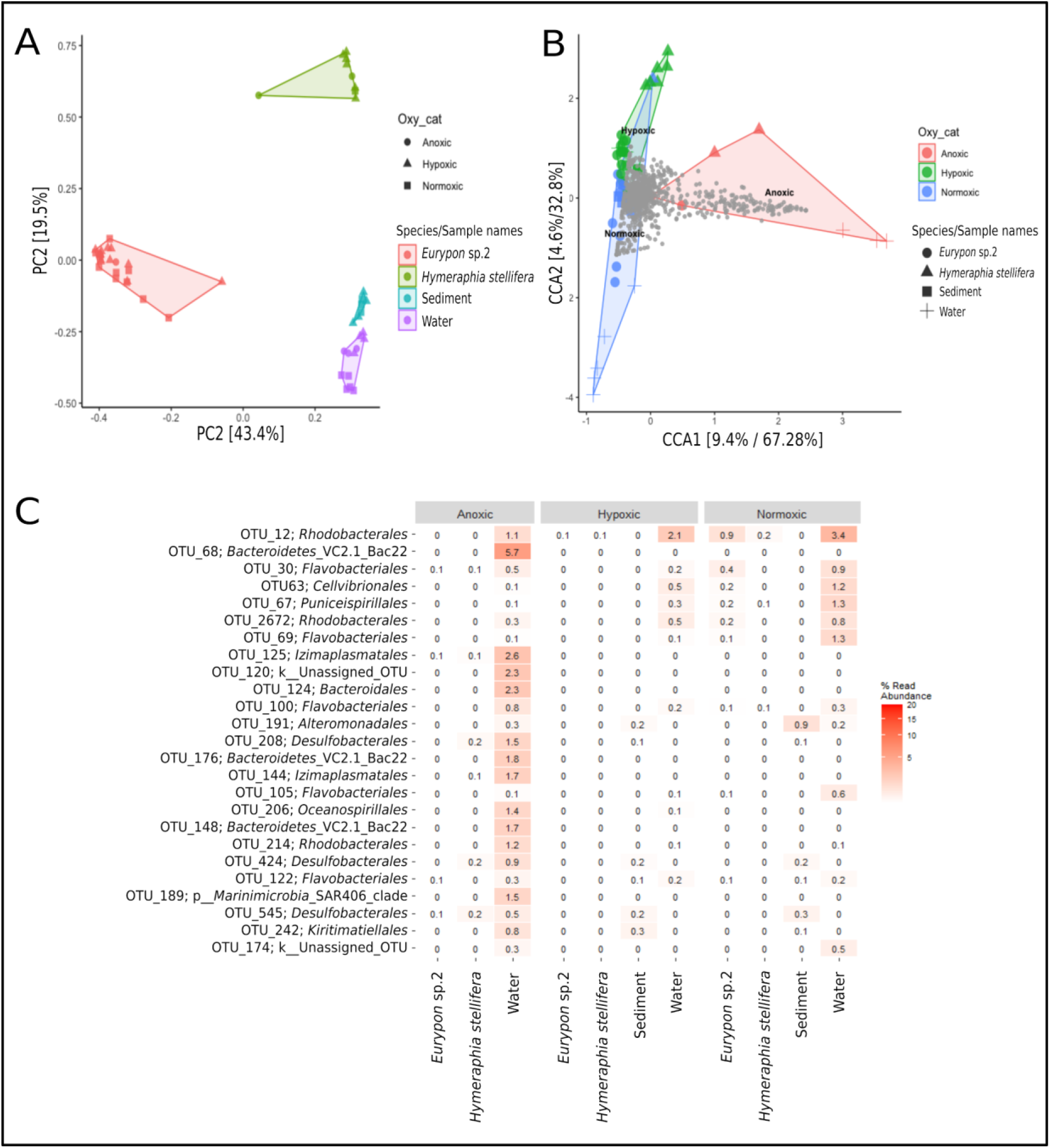
A. PCA of the focused subset. B. CCA of the focused subset. Individual OTUs are identified in grey. C. Heatmap of the OTUs (order level) driving the separation based on anoxia in the CCA, i.e. OTUs contained within the red shaded area in B.

In the PCA, samples grouped definitively by sample type, but the different oxygen conditions did not form clear subgroups even within each sample type, except for the community structures in the water samples (Fig. 3A). When the ordination was constrained by oxygen condition in a Canonical Correspondence Analysis (CCA), anoxic samples appeared to separate out more clearly, but this separation was driven by the communities present in the water samples (plus signs in Fig. 3B). The OTUs driving this separation were identified, extracted and plotted in a heatmap (Fig. 3C), further demonstrating that changes in relative abundance within the water microbial community were not reflected in the microbiomes of either sponge species or within the sediment. Even if water samples were removed from the microbial community analysis, the OTUs driving the separation by oxygen condition were still much more abundant in the water than in sponge samples and thus likely represent low levels of contamination from the water around the sponge and in its canals rather than stable symbioses. Outside of the focused subset, the same pattern of OTUs in the water samples driving separation of microbial communities across oxygen conditions was observed for all Labhra cliff samples (Fig. S5).

The relative abundances of the top seven most abundant OTUs were compared individually across sample type and oxygen condition within the focused subset (Fig. 4, Table S1). The less abundant OTUs (i.e. not top 7) all contributed less than 8% of the relative abundance of OTUs to any one species and did not exhibit notable changes in relative abundance with oxygen condition. In *E*. sp.2, the relative abundance of OTU1, a *Nitrosopumilus*-like *Archaeota* (see Fig. 4) was approximately 40% across all oxygen conditions. From all other sample types OTU1 was absent (relative abundance = 0%), except for *H. stellifera* under anoxia where one of the two individuals of *H. stellifera* appreciably took up OTU1 (relative abundance = 7.64%) under but the other did not (relative abundance = 0.01%). No significant differences were found between relative abundances of OTU1 in *E*. sp.2 under different oxygen conditions (P>0.05, Table S1).

**Fig. 4.**
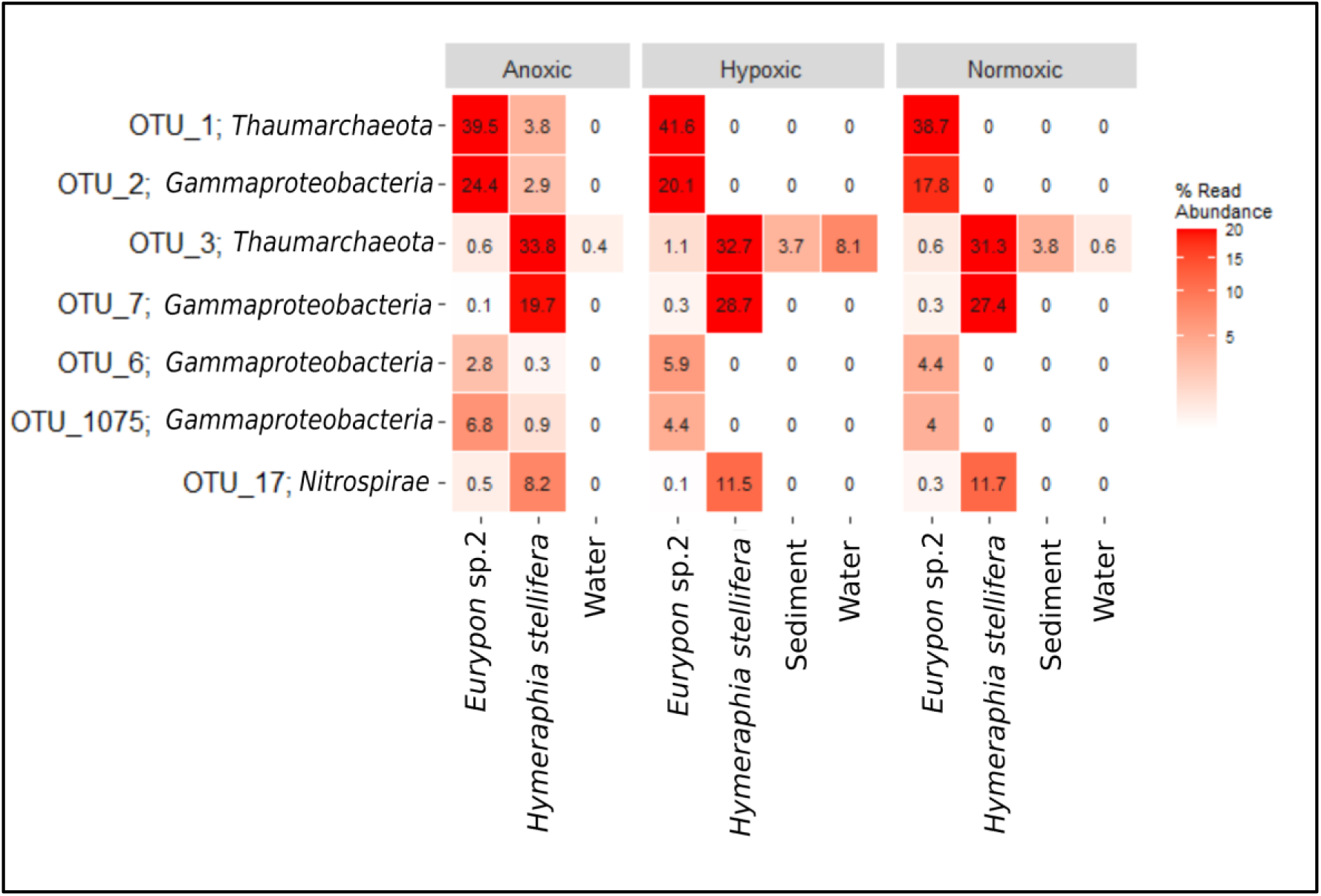
Heatmap of top seven most abundant OTUs in the focused subset.

Similar to OTU1, relative abundances of OTU2, a *Gammaproteobacteria* (see Fig. 6 phylogeny), in *E*. sp.2 were approximately 20% throughout all oxygen conditions, and OTU2 was absent from all other sample types except for *H. stellifera* during anoxia. Within anoxic *H. stellifera*, relative abundances of OTU2 ranged between 5.75 and 0.01% for the two individuals. A Kruskal-Wallis rank sum of only samples from *E*. sp.2 revealed no significant differences in relative abundance of OTU2 between oxygen conditions (P>0.05, Table S1). In contrast, OTU3 and OTU7, a *Nitrosopulmilus* and a *Gammaproteobacteria* (see Fig. 5 and 6 for phylogenies), respectively, were more abundant in *H. stellifera* than in *E*. sp.2.

**Fig. 5.**
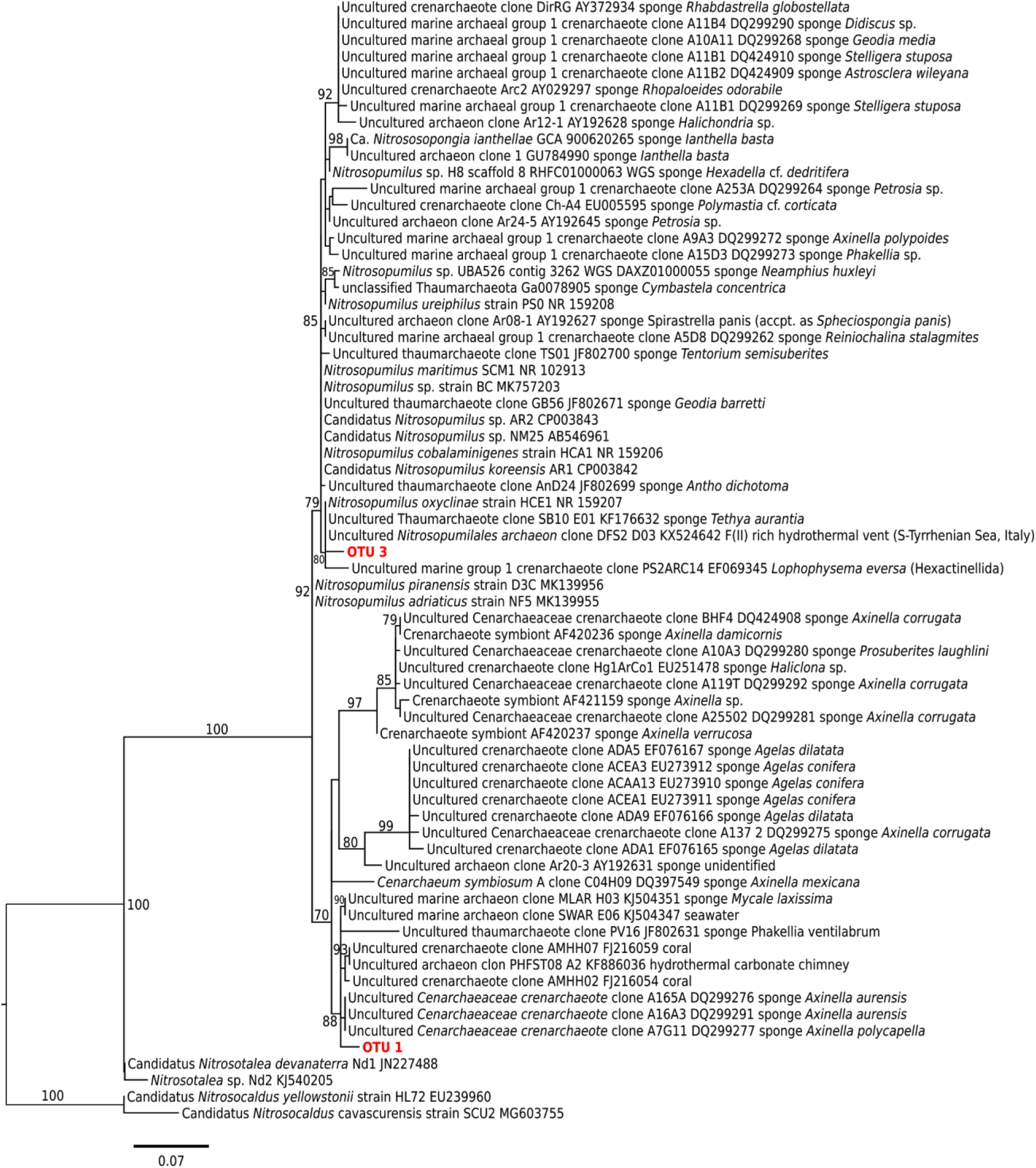
Archaeal Maximum Likelihood phylogeny of partial 16S rRNA gene (292 bp), bootstrap support values (1 000 replicates, GTRCAT model) are given for nodes with 70% or greater. OTUs of interest.

**Fig. 6.**
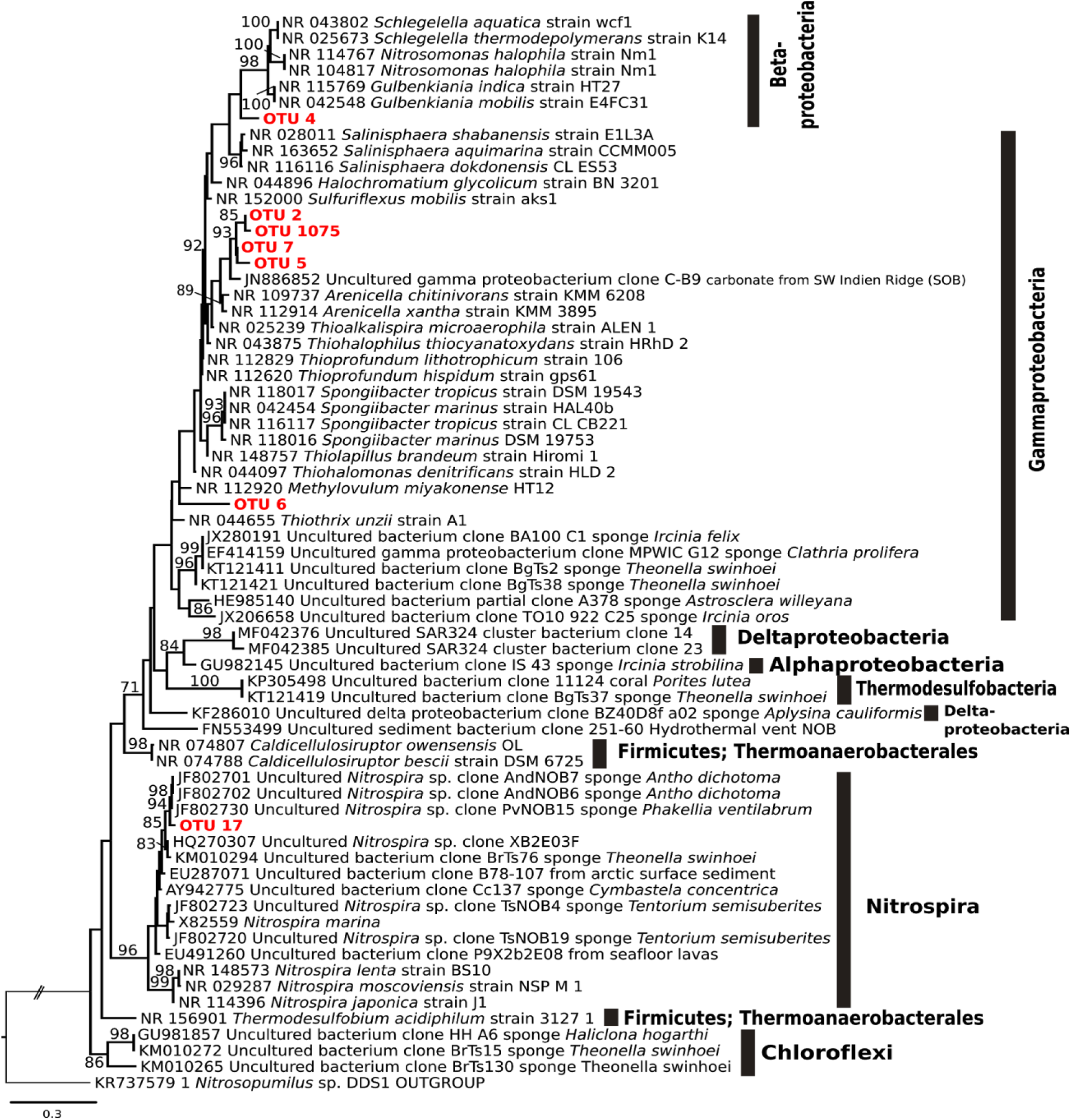
Bacteria Maximum Likelihood phylogeny of partial 16S rRNA gene (292 bp), bootstrap support values (1 000 replicates, GTRCAT model) are reported for nodes with 75% or greater. Top 7 (sponge) OTUs from this study are highlighted in red.

Unlike all other top OTUs (Fig. 4), OTU3 was also found within the sediment and water in addition to sponge samples. Relative abundances in OTU3 were significantly higher in *H. stellifera* compared to any other sample type (P<0.001); however, there were no significant differences between relative abundances of OTU3 within *H. stellifera* under different oxygen conditions (P>0.05, Suppl. Table 1).

Relative abundances of OTU3 within *H. stellifera* were stable at approximately 32% under all oxygen conditions. Relative abundances of OTU3 were significantly lower in *E*. sp.2 than in any other sample type 0.6–1.1% (P<0.05), but no significant changes were observed between oxygen conditions.

Within the water and sediment, average relative abundances of OTU3 ranged from 0.4 to 8.1%, but were not significantly different from one another. The relative abundance of OTU3 was significantly higher in hypoxic waters than normoxic waters (P<0.05, Table S1), but no other significant differences were detected.

Overall, relative abundances in OTU7 were significantly greater in *H. stellifera* than in any other sample type (P<0.001), and these differences held during pairwise comparisons (P<0.001, Table S1). There was a significant decrease in relative abundance of OTU7 in *H. stellifera* under anoxia compared to hypoxia and normoxia (P<0.001, mean relative abundances: 19.7, 28.7 and 27.4%, respectively, Fig. 4). Relative abundances of OTU7 also appeared to decrease within *E*. sp.2 from 0.3 and 0.4% in normoxia and hypoxia to 0.1% in anoxia, but this difference was not significant (P>0.05). The *Gammaproteobacteria* OTU6 and OTU1075 were absent in the sediment and water samples but present in all *E*. sp.2 samples, and only present under anoxia in *H. stellifera*. For *H. stellifera* in anoxia, the relative abundance for OTU6 was 0.549 and 0.002% for both replicates, while the abundance of OTU1075 was 1.89 and 0% for each replicate. No significant changes in the relative abundance of OTU6 were observed in *E*. sp.2 across oxygen conditions. The relative abundances of OTU1075 increased significantly (p<0.05) from approximately 4% in hypoxia and normoxia to approximately 6% in anoxia. The *Nitrospira* OTU17 was also present in both sponge species in all conditions, but relative abundances were significantly higher in *H. stellifera* (∼8–12%) than in *E*. sp.2 (∼0.1–0.5%). No significant changes in relative abundance in OTU17 were observed based on oxygen condition.

Returning to the full dataset (Fig. 7), all sponge species collected from Labhra cliff contained only a few key OTUs that constituted between 30–65% of their microbiomes, depending on the sponge species, and these patterns were generally host-species specific or genera specific (Fig. 7). This trend was also true when relative abundance was examined at the microbial class level (Fig. S6). These key OTUs (OTU1–3, OTU4, 5 and 7) were absent in outgroup samples with the exception of OTU3, which was present in all samples, including the sediment and water, and exhibited relatively high concentrations (5.9 %) in the outgroup species: *Tethya citrina* (Fig. 7). Focusing on only those species verified as present during anoxia, *Raspaciona* sp. contained high levels of OTU3 (*Nitrosopumilus*-like) and OTU10 (unassigned), which were maintained during anoxia and normoxia (Fig. 7). *Mycale* sp. shared the same key symbiont OTUs with *E*. sp.2, but this was the only case where different sponge genera had the same top two symbionts at similar relative abundances. *Mycale* sp. also contained a *Gammaproteobacteria* (OTU27) at relative abundances of 7%, which were absent in *E*. sp.2.

**Fig. 7.**
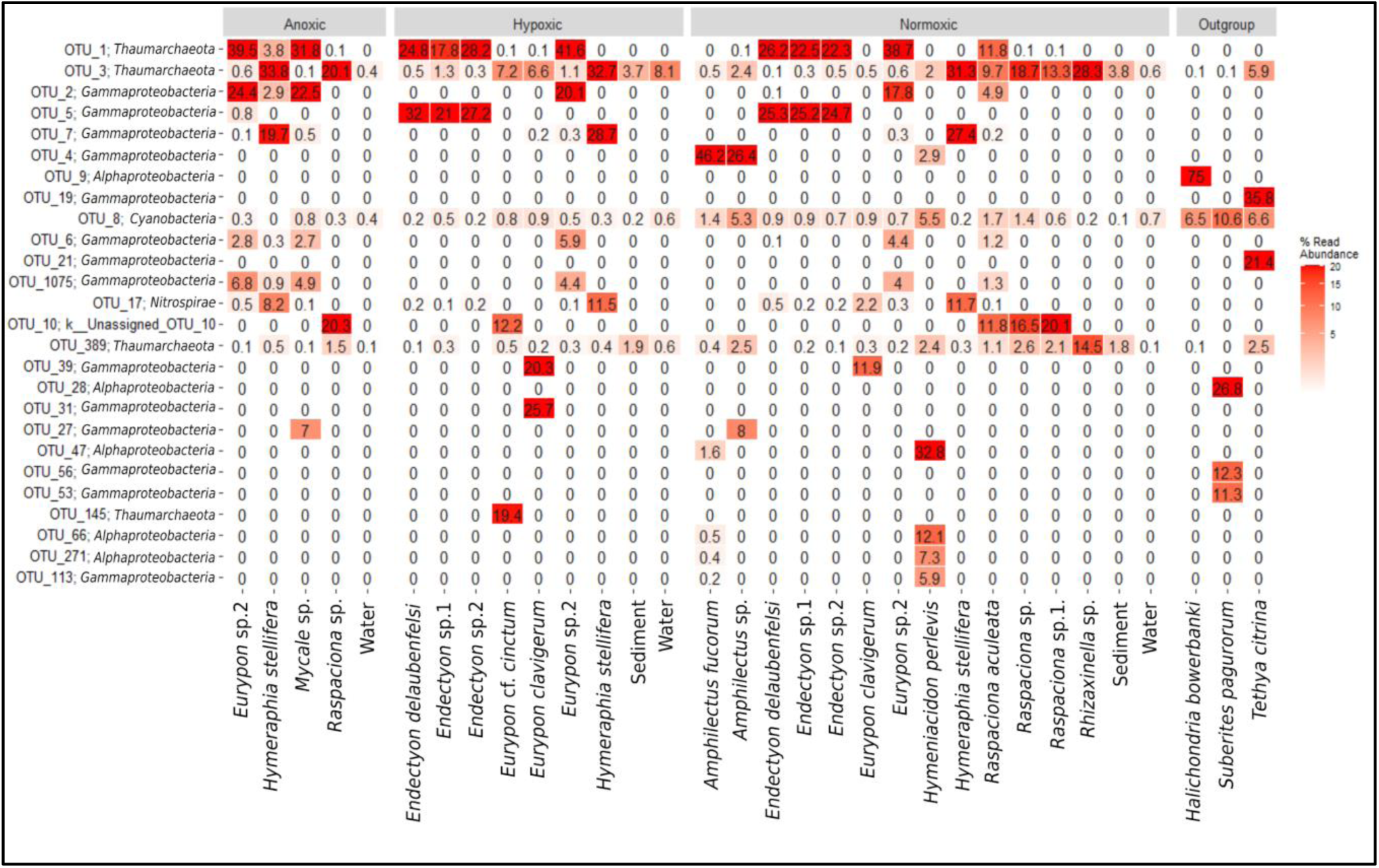
The top 26 most abundant OTUs present in sponge species and environmental samples from the location of seasonal anoxia (i.e. Labhra cliff, designated as anoxic, hypoxic or normoxic) and outgroup samples).

## Discussion

A high level of cryptic sponge diversity was identified at Labhra cliff within sponges that were visually identical *in situ*. Within these sponges, microbiomes were stable under extreme changes in ambient oxygen concentrations despite notable changes in the water-column microbial communities. Four holobiont species that clearly survived prolonged water-column anoxia exhibited three distinct symbiont combinations, all of which were characterised by large populations of *Thaumarchaeota* and either a *Gammaproteobacteria* or an unidentified OTU. These combinations were mostly host species-specific, but some notable symbiont sharing occurred across host taxa, indicating that some symbiont combinations may be better adapted to anoxia than others. These symbionts may confer anoxic tolerance based on anaerobic metabolisms within their phylogenies (Fig. 5 and Fig. 6), which are discussed below. However, there are also potential adaptations of the sponges themselves to deoxygenation, e.g. reduced metabolic rates, as well as the potential of the holobiont to shut down metabolic activity under anoxia that requires further study (see ‘Potential adaptations of the sponge host to seasonal anoxia’). Regardless of the role of the microbiome in deoxygenation tolerance, our observations of Lough Hyne sponges have important implications for ancient and future marine environments.

### Sponge diversity at a seasonally anoxic site

There was a high level of cryptic diversity/speciation within the orange-red encrusting sponges, and at least eight such species are confirmed here as tolerant to seasonal anoxia and hypoxia (*Eurypon clavigerum, Eurypon* cf. *cinctum, Endectyon* sp.1, *Endectyon* sp.2, *E*. sp.2, *H. stellifera, Mycale* sp., and *Raspaciona* sp.). In a previous study, Bell and Barnes (20) also noted the presence of a whitish encrusting species: *Paratimea constellata* and an arborescent species: *Stelligera rigida* (now *Stelligera montagui* (73)), both Axinellida, at 30 m on Labhra cliff. In the current study, *P. constellate* was observed (but not sampled); however, *S. montagui* was not observed. Since two, unidentified massive sponges were also observed (pers. obs. Strehlow) and ‘other’ unidentified species were observed by Bell and Barnes (20), the actual diversity of sponges at this depth is higher than reported here. Despite this diversity, there was still a marked decrease in the overall abundance and diversity of sponge species in the seasonally anoxic waters below 24 m (20). This observation indicates that sponges below 24 m are specifically adapted to this environment and these adaptations could be reflected in their microbiomes.

### Potential adaptations of the microbiome to seasonal anoxia

In Labhra Cliff sponges, microbiomes were largely stable within individual species under different oxygen conditions, despite substantial changes in microbial populations in the water column, suggesting possible adaptations to deoxygenation within the microbiome. Similar stability of sponge microbiomes have also been noted in other studies where sponges have been exposed to sediment loading (36), thermal stress (74), and food shortage (75). Sponge microbiomes are only significantly disrupted when sponges experience physiological stress such as bleaching (76, 77), necrosis (34, 78), disease (79) or mortality (36). Therefore, the stability in the microbiomes we observed in Labhra Cliff sponges could indicate that the holobionts were ‘healthy’ and adapted to periods of anoxia.

Although microbiomes were mostly host species-specific across the whole dataset (Fig. 7), some symbiont strategies were shared across sponge taxa in anoxic-tolerant species. In these four sponge species (*Eurypon* sp. 2, *H. stellifera, Mycale* sp., *Raspaciona* sp.; Fig. 7), three symbiont combinations were identified. These combinations were characterised by their most abundant OTUs as follows: i) OTU1 and OTU2, ii) OTU3 and OTU7, and iii) OTU3 and OTU10 (Fig. 4 and Fig. 7). Combination i) was exhibited by *E*. sp.2 and *Mycale* sp., and *H. stellifera* and *Raspaciona* sp. had combinations ii) and iii), respectively. All combinations included high abundances of *Thaumarchaeota* (either OTU1 or OTU3). Combinations i) and ii) are also dominated by large populations of the *Gammaproteobacteria* OTU2 and OTU7, respectively. In addition to the *Thaumarchaeota* OTU3, combination iii) was characterized by high abundances of the unknown OTU10.

While *Gammaproteobacteria* and *Thaumarchaeota* are common sponge symbionts and often co-occur in one host (40), the combinations of these specific OTUs at their high relative abundances may be unique to anoxic-tolerant species (Fig. 5, 6). There was strong evidence of convergence towards combination i), since it was only acquired by *H. stellifera* under anoxia and was shared across a large host phylogenetic distance, i.e. between the poecilosclerid *Mycale* sp. and the axinellid *E*. sp.2. Although many emergent properties of sponge microbial communities, e.g. community complexity and interactions, are conserved across porifera, it is rare that specific OTUs are shared across large host phylogenetic distances (40). This exceptional symbiont commonality as well as the acquisition of combination i) by *H. stellifera* indicated that this combination may be better adapted to seasonal anoxia than combination ii). Combination iii), conversely, may represent a strategy just as successful as combination i), given its stability in anoxia within *Raspaciona* sp. (Fig. 7).

Both *Thaumarcheota* OTUs were part of the *Nitrosopulmilacae* family, but OTU1 is present only in sponges, making it ‘sponge-specific’; whereas OTU3 was also present in sediment and water samples, making it a generalist. The OTU3 is part of a clade that contains more free-living *Thaumarchaeota* members, including *Nitrosopumilus maritimus* (Fig. 5), than symbionts. Conversely, OTU1 forms a clade that is almost exclusively sponge or coral associated (Fig. 5). Based on the genomes of their close relatives, both OTU1 and OTU3 are likely AOAs and could therefore oxidize sponge-derived ammonia, detoxifying the holobiont and potentially providing a carbon source for the host via chemolithotrophic carbon fixation (80, 81), making them integral parts of holobiont metabolism.

Both ammonia oxidation rates and carbon fixation rates by an AOA symbiont are positively correlated within the sponge *Ianthella basta* (44). Similarly, *Thaumarchaeota* are the main drivers of nitrification in four cold-water sponges (41 a, 42 b). The AOA symbionts of a glass sponge living in mild hypoxia also possess elements of a facultatively anaerobic metabolism, including fermentation and fumarate, nitrate, and sulfite respiration (19). The microbes themselves could also be a food source for the sponge (49, 82, 83). Thus, it is possible that hypoxic environmental conditions are beneficial for the holobiont given the low-oxygen requirements of sponges (16, 17) and the high abundance of *N. maritimus* in marine OMZs (84). Accordingly, the relative abundance of OTU3 significantly increased in the water (but not in any sponge species) during hypoxia but was not significantly different between anoxia and normoxia. Despite this increased abundance in the environment under hypoxia, populations of OTU3 were not significantly increased in *E*. sp.2 under the same conditions. Hypoxia, however, is not the ‘typical’ condition between 25–30 m during the summer in Lough Hyne; instead, anoxia is typical in summer (68). Assuming the sponges are active and pumping (see next section) and given that ammonium oxidation requires oxygen in *Archaea* (48), holobiont metabolisms may be very different under anoxia. Furthermore, *Thaumarchaeota* are functionally diverse (e.g. 44), so the actual metabolisms and symbiotic functions of *Thaumarchaeota* in Lough Hyne sponges need to be verified under their respective conditions.

Like *Thaumarcheota, Gammaproteobacteria* symbionts may contribute key functions to their holobionts, including some that provide tolerance to deoxygenation. Although no single *Gammaproteobacteria* OTU occurred in relative abundances greater than 5.2% in *Raspaciona* sp., symbiont combinations i) and ii) contain high relative abundances of the *Gammaproteobacteria* OTU2 and OTU7, respectively. Unlike the *Thaumarchaeota*, both OTU2 and OTU7 were sponge-specific. Within the focus taxa dataset (Fig. 4), the significant decrease in OTU7 in *H. stellifera* during anoxia, compared to other oxygen conditions, may correspond to the appearance of OTU2, if both occupy the same niche. The same might be true of another *Gammaproteobacteria*, OTU1075, which significantly increased in relative abundance in anoxia in *E*. sp.2 and was more closely related to OTU2 than OTU7 (Fig. 6). The facultative anaerobe *Thioalkalisira microaerophila*, which can use sulfide as an election donor and grows in micro-oxic conditions (85), as well as *Thiohalophilis thiocyanatoxydans*, which can grow anaerobically using thiosulfate as an electron donor and nitrite as an electron acceptor (85), are in the same clade as the sponge-specific *Gammaproteobacteria* (Fig. 6). Therefore, it is possible that the sponge-specific *Gammaproteobacteria* in our samples possessed both aerobic and anaerobic capacities.

In addition to the two most abundant OTUs in combination ii), a *Nitrospira* (OTU17) was found in high relative abundances in *H. stellifera*. Although it was absent or significantly less abundant in other Labhra cliff sponges (Fig. 7), OTU17 may perform important metabolic functions within the *H. stellifera* holobiont. The OTU17 is closely related to a *Nitrospira* (CcNi) that is associated with the sponge *Cymbastela concentrica*, and even though some *Nitrospira* can completely oxidize ammonia to nitrate (commamox), OTU17 may only oxidize ammonium to nitrite as was predicted for CcNi (86). Although it was not significant, relative abundances of OTU17 decreased during anoxia in *H. stellifera*, which could be due to a lack of oxygen inhibiting the metabolism and growth of *Nitrospira* species. Although *Nitrospira* are also conspicuously absent (see also Figure 3, 40, 44) or inactive (42) in some sponge holobionts in general, CcNi symbionts in *C. concentrica* form close metabolic associations with the host and other microbes, including a *Thaumarchaeota*: CcThau (86).

Coincidentally, CcThau is more closely related to OTU3 than to OTU1 (Fig. 5), and therefore the co-occurrence of relatively large populations of OTU3 and OTU17, could indicate a co-evolution between these two OTUs. The acquisition of OTU1 by *H. stellifera* could therefore disrupt these partnerships under anoxia (87), but this remains to be tested. It is also possible that OTU17, like some of its congenerics, could perform comammox and/or hydrogen oxidation coupled to sulfur reduction in anaerobic conditions, making it well adapted for low-oxygen stress (88). In either case, the *H. stellifera* holobiont likely employed a separate metabolic strategy under normoxia compared to anoxia and when compared to hosts with symbiont combination i) or iii).

The holobionts from the genera *Raspaciona* likely employ different metabolic strategies in response to anoxia. Like *H. stellifera*, they host large, stable populations of OTU3, but *Raspaciona* sp. does not acquire more OTU1 or any OTU2 populations in anoxia. Instead, *Raspaciona* sp. harbored large, stable populations of the unidentified OTU10 (Fig. 7), which was completely absent from all other samples except for one *Eurypon* cf. *cinctum*, taken under hypoxia. The unassigned OTU10 has an unknown metabolism; however, it’s stability in *Raspaciona* sp. through anoxia and normoxia, and its presence in *E*. cf. *cinctum*, indicates that it may confer some degree of deoxygenation tolerance.

Microbes associated with sulfate reduction and anammox were conspicuously absent or present in very low abundances in Labhra Cliff sponges. Probable sulfate reducing OTUs were present in the anoxic water at much higher abundances than in any sponge species under the same conditions, but they were absent in all sponges under normoxic and hypoxic conditions (Fig. 3C). The *Planctomycetes* as a phylum, which contains anammox bacteria, were present at low levels in all samples and paradoxically appear to decrease in relative abundances in anoxic water in *E*. sp.2 and *H. stellifera* (Fig. S6). In *Raspaciona* sp., conversely, the relative abundance of this phylum increases in anoxia from 2.4% to 4.7% (Fig. S6). With the exception of *Raspaciona* sp., the low signals of *Planctomycetes* in the other sponges were likely contamination from microbes in the sponge water canal system and are usually bioinformatically filtered out of analyses of symbionts (e.g. 89). Curiously, both sulfate reduction and anammox bacteria have been confirmed in the holobiont *G. barretti*, which experiences internal anoxia in its tissues (49, 50). The difference in the microbial communities in *G. barretti* compared to anoxic-tolerant species from Lough Hyne could be due to differences in morphology as *G. barretti* is a massive species, or environment, since it is found under more or less constant oxygen rather than seasonal anoxia. Thus, the ‘anoxic micro-ecosystems’ observed in *G. barretti (49)* may result from its morphology, and the thin, encrusting sponges of Lough Hyne could be comparatively more oxygenated most of the year, even if pumping ceases (90).

Periods of pervasive oxygenation would restrict symbioses with obligate anaerobes and favor microbes with flexible metabolic strategies.

Notwithstanding these potential anaerobic processes, the three symbiont combinations outlined above do not universally confer hypoxic tolerance to sponges in general. For example, *Thaumarchaeota* are effectively absent in the hypoxic-tolerant species *H. panicea* (91). Moreover, *H. panicea* contains high abundances (>75%) of an *Alphaproteobacteria* as did its congeneric *H. bowerbankii* (Fig. 7), and *Alphaproteobacteria* were effectively absent from anoxic-tolerant Lough Hyne sponges. Although the microbial community within the hypoxic-tolerant *T. wilhelma* has not yet been investigated in detail, only two bacterial genomes were identified from genomic sequencing of *T. wilhelma*, and both were likely *Alphaproteobacteria* (92). It is therefore unclear if *T. wilhema* contains *Thaumarcheota* OTUs in high abundance, though it’s congeneric, *T. crypta*, does (Fig. 7). An *Alphaproteobacteria* cultured from the Lough Hyne sponge *Axinella dissimilis* was able to grow anaerobically via fermentation and denitrification (93). Therefore, many microbiome structures may be capable of coping with hypoxia. Nevertheless, neither the hypoxic tolerant *H. panicea* (17) nor *T. wilhelma* (16) tolerate prolonged anoxia, so the Labhra cliff holobionts may be uniquely tolerant to anoxia even compared to other poriferans. These sponge hosts may also be adapted to tolerate deoxygenation directly, or the ability to survive anoxia may depend on metabolic shutdown of either the host, symbionts or both.

### Potential adaptations of the sponge host to seasonal anoxia

It is unlikely that Lough Hyne sponges die off *en masse* during anoxia and recolonize the area during normoxia because no dead tissue or discolouration was present around anoxic sponges. It is possible that these sponges decrease or cease metabolic activity during anoxia (25, 94), but this requires further investigation. For some marine invertebrates, environmental anoxia can trigger a switch to a fermentation-based metabolism, which results in a considerably decreased metabolic rate; however, the byproducts of fermentation still need to be eliminated into the environment (reviewed in 95). If this elimination cannot be achieved by diffusion alone, it is possible that the sponges continue to pump during anoxia albeit at a potentially decreased rate, and thereby still provide dissolved organics, ammonia, CO_2_, and other metabolic products to their symbionts.

Nonetheless, Lahra Cliff sponges were definitely pumping under hypoxic conditions (Fig. 1A), which is consistent with normal transcription activity in *T. wilhelma* under hypoxia (16) and observations of sponges inhabiting consistently hypoxic environments (96, 97). Pumping was unsurprising considering that mobile fish and crabs were also observed under hypoxia at Lough Hyne (pers. obs. Strehlow), and oxygen levels were above lethal and sublethal thresholds of many fish and invertebrates (12). Nevertheless, in a separate study, a single individual of *G. barretti* drastically reduced its pumping rate following *ex situ* oxygen depletion (20% air saturation) (98), and feeding rates in one individual (out of three) of *H. panicea* were reduced in low-oxygen concentrations (3% air saturation) (17). The sublethal, physiological impacts of deoxygenation, therefore, also need to be considered in the future.

Despite deoxygenation tolerance in some species, overall, sponge diversity and abundance is likely limited by seasonal anoxia (20). Growth and reproduction may be impacted by seasonal anoxia because collagen synthesis is oxygen dependent; however it is still possible in very low oxygen concentrations (see 99). Sponge larvae may also require elevated oxygen due to their motility. Also, elevated oxygen may be needed during settlement and early development in sponges; however, the specific oxygen requirements for these life stages remain unknown. Moreover, a combination of factors could restrict settlement and growth to fewer species. Although sedimentation rates in the anoxic region are equivalent to that of other sites in Lough Hyne (100), the combination of sediment and anoxic stress could restrict sponge distributions. Additionally, seasonally decreased metabolic activity in the holobiont could limit growth or the production of secondary metabolites under anoxia, leading to increased spongivory in normoxic months when mobile predators return.

It is also possible that, like the sponges, some or all of the microbiome could become dormant under anoxia. The capacity to become dormant is common and phylogenetically widespread in microorganisms (reviewed in 101) and may occur in response to environmental stress including hypoxia (102). For instance, pelagic *Thaumarchaeota* are present in sulfidic zones, but they exhibit lower expression levels for genes involved in ammonia oxidation and may be inactive (103).

However, as stated in the previous section, *Thaumarchaeota* in symbiosis with sponges may possess elements of anaerobic metabolism (19) that could aid the sponge holobionts under anoxia if they are active. The major *Gammaproteobacteria* OTUs could similarly be inactive during anoxia in Lough Hyne, though there is also strong evidence of anaerobic metabolisms within the clade formed by these OTUs within the anoxic tolerant sponge holobionts. Moreover, even dormant microbes require some maintenance of proton motive force (102) or DNA repair (104). So there may be some activity within a dormant microbiome under anoxia, which could perhaps be linked to the host’s decreased pumping rate suggested above for waste elimination. Finally, if microbial dormancy occurs, it could still convey some resilience to the holobiont, stabilizing the microbiome under deoxygenation stress and allowing for rapid recovery following reoxygenation.

### Implications for early animal evolution and future oceans

These results have important implications for early animal evolution and the state of future oceans. Under the lower oxygen concentrations of the Neoproterozoic Era, early metazoans likely tolerated both widespread hypoxia and transient anoxia (105–107). The Labhra Cliff sponges demonstrated that this ancient tolerance could have involved the microbiome, but how analogous are these holobionts to early metazoans? The ancestral sponge evolved in a microbial world and may have consequently formed close associations with many symbionts like modern sponges. Modern symbioses with both *Gammaproteobacteria* and *Thaumarchaeota* were present in all 81 species assessed by Thomas et al. 2016 (40). It is therefore conceivable that the ancestral sponge contained either *Gammaproteobacteria, Thaumarchaeota*, both, or their respective ancestral forms. The loss or decreased abundances in these symbiont groups in sponge lineages that evolved into our ‘outgroup’ samples (Fig. 7) could then correspond with the absence of this group in the seasonally anoxic site. Since Lough Hyne is geologically young (69), its sponges may not have had time to evolve adaptations to its seasonal anoxia, hinting at innate abilities within these sponges that could be holdovers from the Neoproterozoic Era. Nevertheless, faster processes could be at play, e.g. transgenerational acclimation (108) or the horizontal acquisition of tolerant symbionts, permitting ‘rapid’ colonization of the seasonally anoxic niche.

Additionally, If encrusting sponges are especially tolerant to fluctuating redox conditions, then early sponges in the Cryogenian Period (720–635 Myr) may have also been thin, encrusting, and rare and thus less likely to fossilize, which explains gaps in the fossil record and its disagreement with molecular clock results (see also 109). Consequently, the first metazoans could have been very similar to the Labhra cliff sponges in their symbiont composition and morphology, and even if it is only through convergent evolution, the seasonally anoxic sponges of Lough Hyne are an important model system for studying early animal evolution.

Considering the past also yields clues about the future. Some previous mass extinction events were probably driven by acidification, warming and deoxygenation of the oceans following extensive volcanism (12, 110–113). Since similar stressors are facing modern oceans as a result of anthropogenic CO_2_ release, could sponges outcompete corals in future scenarios (e.g 114, 115)?

Sponge abundance has recently increased on some coral reefs, due in part to a decrease in coral abundance (116–118), but the future may be more nuanced. Recent experiments showed that the necrosis and bleaching caused by thermal stress was ameliorated by increased CO_2_ in two phototrophic sponge species under scenarios equivalent to the worst-case warming predictions, i.e. RCP8.5 (119), but necrosis was exacerbated by these two stressors in two heterotrophic species (120). Moreover, not all phototrophic sponges have this advantage, since at least one species died under these conditions (121), and most experiments do not include oxygen as a factor, which is predicted to decrease by up to 3.7% under RCP8.5 (122). According to a meta analysis across marine benthic organisms, the combination of thermal stress and deoxygenation reduced survival times by 74%, compared to each stressor in isolation, and increased the lethal concentration of oxygen by 16% on average (123). As with increased temperature and CO_2_, responses to deoxygenation are likely species specific as suggested by this study. Some coral (124, 125) and sponge (11) species may be tolerant to deoxygenation while others are not. The effects of the combination of all three stressors, however, are virtually unknown, and thus require extensive research in the future.

## Conclusions

This study demonstrated that distantly related sponge species can tolerate seasonal anoxia, and the cryptic diversity of sponges tolerating these extremes in Lough Hyne is high. Microbiomes of all sponge species were remarkably stable under varied oxygen conditions and mostly species specific, though some OTUs exhibited minor host sharing under anoxic conditions. Three different symbiont combinations were found to correspond with anoxic tolerance in Lough Hyne sponges, but future research is needed to verify the hypothesized metabolic interactions of host and symbionts under anoxia and to untangle microbial community structure from other shared characteristics. These microbial communities are not present in all sponge species with a reported hypoxic tolerance, but these combinations may be crucial for anoxic tolerance if only for their possible capacity for dormancy. The lack of significant disruption of the microbiomes during anoxia indicated that these sponge holobionts are well adapted to anoxia even without the HIF pathway. Similar sponge-microbial associations could have been important in the early evolution of animals under the lower oxygen conditions of the Proterozoic. Although our study suggests some strategies for holobiont deoxygenation tolerance, more research is needed to understand the individual and combined effects of acidification, warming and deoxygenation to predict the structure of future ecosystems and to determine the cause-effect pathways of these stressors in order to inform management and restoration efforts.

## Materials (or Subjects) and Methods

### Study site

Sampling was performed in the Lough Hyne Nature Reserve (51°29’N, 09°18’W), a semi-enclosed marine lough in County Cork, Ireland. Lough Hyne is connected to the Atlantic Ocean by a narrow (∼25 m), shallow tidal rapid. Most samples were taken from Labra Cliff (N 51° 30.0530’ W9° 18.1767’), located in an area known as the Western Trough (Fig. 1A). The water current speed at this site is < 5 cm^−1^, and the cliff surface, where sponges were found, is covered with a thick layer of silt (20). A seasonal thermocline exists and develops at ∼25 m when surface temperatures increase in summer months. Below this depth an anoxic/hypoxic layer forms (68).

### Physical data

Oxygen concentrations and temperature were measured at increasing depths using a Pro20 Dissolved Oxygen Meter (YSI, USA) (Fig. 1). Anoxic, hypoxic and normoxic conditions were defined by dissolved oxygen concentrations of 0.00–0.01 (i.e. the detection limit of the instrument), 1.30– 3.56, and 5.3–12 mg L^−1^, respectively. A conductivity, temperature and depth (CTD 90, Sea and Sun Technology, Germany) instrument, equipped with high range (0.2–300 μM) and low range (0–200 nM) oxygen sensors, was used to more accurately measure low concentrations of oxygen in July 2019. Temperature, turbidity and salinity values measured from the CTD during July 2019 are reported in Fig. S1, while photosynthetically active radiation (PAR) and chlorophyll concentrations measured by the CTD are reported in Fig. S2 (A,B). The concentrations of photosynthetic pigments (Chlorophyll *a* and pheophytin) were measured in July 2019 from water samples taken from 0–35 m at a distance of 5 m and at 39 m using a battery powered pump (see 126) and description in Supplementary material (Fig. S2C).

### Sampling and DNA extraction

Sponges, water and sediment were sampled at Labhra Cliff using SCUBA between July 2018 and August 2019 under permit no. R23-27/2018 issued by the Irish Department of Environment, Heritage, and Local Governments. A two-point calibrated HOBO Dissolved Oxygen Logger (U26-001, Onset, USA) was used during all dives to measure dissolved oxygen (mg/L) and ensure that all sampled sponges were assigned to the correct oxygen condition, i.e. anoxic, hypoxic or normoxic. Reference sponge samples were also taken above the 24 m depth, where oxygen is continuously present. The collection depths were recorded, and the oxygen condition of these samples was noted as ‘normoxic’ (Fig. 2 and Fig. S3 for full sampling details). Sulfide concentrations were measured from two 50 mL water samples at 27 m at Labhra cliff and flash frozen in liquid nitrogen. After freezing, water samples were post-fixed in 5% zinc acetate, and sulfide concentrations were measured spectrophotometrically following: (127). The freezing may have degassed the samples and some sulfide may have oxidized after the samples were collected but before they were frozen. Sponges from the Labhra cliff site were sampled using a flathead screwdriver to scrape off a ribbon of tissue that was then removed from the encrusted rock and placed into a whirl-pak bag using forceps. This process was optimized to both use minimal movement, limiting sediment resuspension, and ensure that divers stayed within no-decompression limits. Still, the depth and low visibility restricted the number of samples that could be taken, particularly during the anoxic period. All samples were frozen in liquid nitrogen soon after divers reached the surface (a few minutes). Before sampling, the pumping status of five individual sponges under hypoxic conditions, and five individuals under normoxic conditions was assessed visually using fluorescein dye. This procedure was not possible during anoxia due to time and visibility constraints.

Genomic DNA was isolated from sponge tissue (anoxic = 6, hypoxic = 25, normoxic = 39, total = 70) using the DNeasy (Qiagen, Germany) Blood and Tissue Kit protocol. Sediment samples (hypoxic = 4, oxic = 3, total = 7) were extracted using the DNeasy PowerSoil Pro Kit (Qiagen, Germany). Sterivex^™^ Filters (Millipore, Billerica, MA, USA) were used to filter one liter of seawater (anoxic = 3, hypoxic = 4, oxic = 5, total = 12), and the protocol from (128) was used to open filter casings and a standard phenol-chloroform protocol (129) was performed to extract genomic DNA.

For microbial community comparison, ‘outgroup’ samples were taken from three different species: *Halichondria bowerbanki, Suberities pagurourm* and *Tethya citrina* (n=3 per species) from a separate location (51.499877, -9.296661, Fig. 1A) at a depth of 2 m. These species were not observed at the seasonally anoxic site. For all sponge samples sampled from Labhra cliff, less than 5% of the sponge volume was sampled, leaving sufficient biomass for survival and recovery (130).

### Sponge identification

All sponge samples were identified to genus and species level according to their phylogenetic position relative to known species by amplification of the *cox1*, (∼659 bp) gene using primers dgLCO1490 and dgHCO2198 (131), and the 28S (C-region, ∼550 bp) gene using primers C2 and D2 (132). Amplifications and clean-up followed the protocol in (133). The remaining supernatant was sequenced by Macrogen (Europe). BLAST searches against NCBI GenBank (https://blast.ncbi.nlm.nih.gov/Blast.cgi) were used to confirm sponge origin. Raw trace files were processed in Geneious® v.11.1.5 (https://www.geneious.com). Alignments were generated separately for each gene using the MAFFT plugin v.1.4.0 in Geneious® with the L-INS-I algorithm (134), due to the heterogeneous taxon sampling and moderate sequencing success of *cox1*. Phylogenetic tree reconstructions were performed using the MrBayes 3.2.2 plugin on the CIPRES Science Gateway (135). The best fit evolutionary model (GTR+G+I) was selected according to the results of JModelTest2 (136) plugin using the CIPRES Science Gateway v.3.3 (135). Two concurrent runs of four Metropolis-coupled Markov-chains Monte Carlo (MCMCMC) for 100 000 000 generations were run and stopped when the average standard deviation of split frequencies was below 0.01. The first 25% of the sampled trees was removed as burn-in for further analyses. FigTree v1.4.2 (137) was used to visualize the trees.

## Microbial community

### Library preparation

Sequencing libraries of *Bacteria* and *Archaea* 16S V4 rRNA genes were prepared based on a custom Illumina protocol (138) at DNASense ApS (Aalborg, Denmark). A detailed protocol for library preparation and purification is provided in the supplementary material. Forward and reverse tailed primers were designed according to Illumina (2015) targeting *Bacteria* and *Archaea* 16S V4 rRNA gene: (abV4-C-f, GTGYCAGCMGCCGCGGTAA and abV4-C-r, GGACTACNVGGGTWTCTAAT) (139, 140). Purified sequencing libraries were pooled in equimolar concentrations and diluted to 2 nM. Samples were paired-end sequenced (2×300 bp) on a MiSeq (Illumina, USA) at DNASense ApS using a MiSeq Reagent kit v3 (Illumina, USA) following the standard protocol. Forward and reverse reads were trimmed and de-replicated, and processed reads were divided into OTUs and assigned relative abundances and taxonomies as outlined in the Supplemental material. The results were analysed in R v. 3.5.2 (141) through the RStudio GUI (142), using the ampvis2 package v.2.5.8 (143). All further analyses were performed using this package or base R unless otherwise indicated.

### Statistical analysis

Samples from the sponges *Hymeraphia stellifera* (anoxic = 2, hypoxic = 6, oxic = 1, total = 9) and *Eurypon* sp.2 (anoxic = 2, hypoxic = 12, oxic = 13, total = 27) as well as from water and sediment were subsetted due to their representative sampling under all three different oxygen conditions to form the ‘focused subset’. Variation in OTU composition within the focused subset was assessed by Principal Component Analysis (PCA) with Bray-Curtis dissimilarities. A Canonical Correspondence Analysis (CCA) of OTU data constrained to oxygen condition, using the Pearson chi-squared measure and “Hellinger” transformation, was also performed on the focused subset to test for any influence of oxygen condition on microbial community structure. A CCA was also performed on all samples taken at Labhra cliff (n=89) as above.

The presence of significant variation within the microbial communities based on sample type and oxygen condition within the focused subset was tested using a 2-factor PERMANOVA with 10,000 permutations with the vegan package (144) in R. Significant differences in relative abundances were also tested between sample type and oxygen condition for the top seven most abundant OTUs individually within the focused subset, using analyses of variance (ANOVAs) for OTUs with normal distributions and Kruskal-Wallis rank sum tests for those with non-parametric data. Tukey Honest Significant Difference (HSD) tests and Wilcox pairwise tests were used for post hoc tests of parametric and nonparametric data, respectively. If one sample type across all treatments had zero percent abundance for a specific OTU, those data were excluded from analysis to meet assumptions of normality.

Separate Bacterial and Archaeal Maximum Likelihood phylogenies of partial 16S rRNA gene were calculated using RAxML v.8 (145) on the CIPRES Science Gateway V. 3.3 (135). Alignments including the ‘focused subset’ of the top bacterial OTUs (OTU4, OTU1075, OTU2, OTU5, OTU7, OTU6 and OTU17) and archaeal OTUs (OTU1 and OTU3) were merged with datasets of Feng et al. 2016 (43).

## Data availability

All sample metadata, OTU table, alignments, physical data and scripts (ampvis2) for this publication are available on github (https://github.com/astridschuster/Microbiome_MS_2020.git). The 16S sequence data are available under the BioProject number XXXX in NCBI. Sponge barcode sequences are deposited at the European Nucleotide Archive (ENA) under Primary Accession Number: PRJEB38772 with 28S accession numbers LR880974–LR881068 and *cox1* LR880934–LR880963 respectively.

## Acknowledgements

Field work at Lough Hyne was carried out with the permission of the National Parks and Wildlife Service of Ireland. This project was funded by VILLUM FONDEN [Grant No. 16518]. AS was partly funded by the H2020 Project SponGES (Grant Agreement no. 679849). The authors would like to thank Luke Harman, Allen Whittaker of University College Cork. We also thank the NordCEE lab technicians of the Department of Biology at SDU for constant support in the laboratory procedures and Laura Bristow for assistance with CTD data and operation. In addition, we thank DNASense for the sequencing of the microbial community and the help with the ampvis2 R package in particular Rasmus Wollenberg and Kasper Skytte Andersen.

## Competing Interests

The authors declare that they have no conflict of interest

## Supplementary material

### Physical data

To determine chlorophyll a (Chl *a*) and pheophytin concentrations, 50 mL of seawater from each depth was filtered through 25 mm diameter pre-combusted (450 °C for 4 h) GF/F filters. The filters were then placed in 2 mL Eppendorf® tubes wrapped in aluminium foil and frozen (−20 °C) until analysis. For analysis, the filters were transferred to 15 mL centrifuge tubes and acetone (8 mL, 90 %) was added. The samples were kept overnight (5 °C) before sonication (30 min) in a sonication bath and then centrifuged (3000 rpm at 6 °C for 5 min). Chlorophyll *a* was measured with a Turner TD-700 fluorometer (Turner Design, Sunnyvale, CA, USA). Samples were then acidified with 50 µL of 1 M HCl and pheophytin was measured. The fluorometer was calibrated using an extract from spinach and serial dilutions of a 4 mg L^−1^ stock standard, 90 % acetone was used as a blank. A solid-state secondary standard (SSS) was measured every ten samples. The SSS insert provides a very stable fluorescent signal and is used when measuring Chl *a* to check for fluorometer stability and sensitivity. The detection limit was 1 μg L^−1^.

### Microbial community sequencing

For library preparations, up to 10 ng of DNA extracted from sponge, water and sediment samples was used as a template for PCR amplification of the 16S V4 rRNA gene amplicons, covering both *Bacteria* and *Archaea*. The PCR reaction (final volume 25 µl) was prepared with the following reagents: dNTPs (100 µM of each), MgSO4 (1.5 mM), Platinum Taq DNA polymerase HF (0.5 U/reaction), Platinum High Fidelity buffer (1X, Thermo Fisher Scientific, USA) and tailed primer mix (400 nM of each forward and reverse primer). The PCR program ran as follows: 95 °C for 2 min, 30 cycles of: 95 °C for 15 s, 55 °C for 15 s, and 72 °C for 50 s; and final elongation at 72 °C for 5 min. Duplicated PCR reactions were performed and pooled after PCR for each sample. Amplicon libraries were purified by a standard protocol for Agencourt Ampure XP Beads (Beckman Coulter, USA) with a bead ratio of 4:5. DNA was eluted in 25 µl nuclease free water (Qiagen, Germany). A Qubit dsDNA HS Assay kit (Thermo Fisher Scientific, USA) was used to measure DNA concentration. Validation of PCR product size and purity of sequencing libraries was carried out via Gel electrophoresis using Tapstation 220 and D1000/High sensitivity D1000 ScreenTapes (Agilent, USA). Sequencing libraries were prepared from the purified amplicon libraries using a second PCR containing PCRBIO HiFi buffer (1x), PCRBIO HiFI Polymerase (1 U/reaction) (PCR Biosystems, UK), adaptor mix (400 nM of each forward and reverse) and up to 10 ng of amplicon library template. The PCR settings were: 95°C for 2 min; 8 cycles of 95 °C for 20 s, 55 °C for 30 s, 72 °C for 60 s; and final elongation at 72 °C for 5 min. Sequencing libraries were purified again using Agencourt Ampure XP Beads (Beckman Coulter, USA) with a beat ratio 4:5, and DNA was eluted in 25 µl of nuclease-free water (Qiagen, Germany). Product size and purity of a subset of sequencing libraries was validated on a Tapstation 220 and D1000/High sensitivity D1000 ScreenTapes (Agilent, USA). A PhiX control library (>10%) was spiked in to overcome low complexity issues often observed with amplicon samples. After sequencing, Trimmomatic v. 0.32 (146) was used to quality trimm forward and reverse reads with settings: SLIDINGWINDOW: 5:3 and MINLEN: 250. The trimmed forward and reverse reads were merged using FLASH v. 1.2.7 (147) with settings: -m 10 -M 250. Trimmed reads were de-replicated and formatted for the use of the UPARSE workflow (148). Dereplicated reads were clustered by usearch v.7.0.1090 -cluster_otus command with default settings. The OTU abundances were estimated using -usearch_global command with -id 0.97 - maxaccepts 0 -maxrejects 0. Taxonomy was assigned using the RDP classifier (149) implemented in the paralles_assign_taxonomy_rdp.py script in QIIME (150), using -confidence 0.8 and the SILVA database, release 132 (151).

### Microbial communities

The following information pertains to observations of sponges species that were not sampled under anoxia, i.e. either only in normoxia or hypoxia. *R. aculeta* contained four key OTUs: OTU1, OTU2 (*Gammaproteobacteria*), OTU3, and OTU10 (unassigned) in relative abundances of 11.8, 4.9, 9.7, and 11.8%, respectively (Fig. 7), all other OTUs present in *R. aculeata* had relative abundances lower than 4%. Similarly, *Hymeniacidon perlevis* had five OTUs that constituted 65% of its microbiome, including OTU47 (*Aphlaproteobacteria, Terasakiellaceae*) at 32.8%, OTU66 (*Alphaproteobacteria, Novosphingobiu*) at 12.1%, OTU271 (*Alphaproteobacteria, Terasakiellaceae*) at 7.3%, OTU8 (*Cyanobacteria, Synechococcus* CC990) at 6.5%, and OTU113 (*Gammaproteobacteria*) at 5.9% relative abundance. In contrast, *A. fucorum* contained only one key symbiont: OTU4 (*Gammaproteobacteria*), but it accounted for 46.2% of the microbiome by relative abundance. Lower relative abundances of OTU4 were observed in *Amphilectus* sp. (26.4%) along with OTU27 (also a *Gammaproteobacteria*) with 8% relative abundance. However, with the exception of *R. aculeta*, the aforementioned sponge species were collected at depths shallower than those exposed to seasonal anoxia (i.e. <24m). For the other 11 sponge species, there were only two key OTUs. The two key OTUs of *Eurypon* cf. *cinctum*, OTU145 (a *Nitrospira*) and OTU10 (Fig. 7).

*Eurypon clavigerum* contained two *Gammaproteobacteria* OTUs as its main symbionts (OTU31 and 39), and the key symbionts of *Rhizaxinella* sp. were OTU3 (*Nitrosopumilus*-like) and OTU389 (candidatus *Nitrosopumilus*). In the remaining sponge species (i.e. excluding *Eurypon* cf. *cinctum, Eurypon clavigerum, Raspaciona* sp., *Raspaciona* sp.1, and *Rhizaxinella* sp.), these two key OTUs consisted of one *Nitrosopumilus* (or *Nitrosopumilus*-like) and one *Gammaproteobacteria*. The specific combination of *Nitrosopumilus* (or *Nitrosopumilus*-like) and *Gammaproteobacteria* OTUs generally depended largely on the sponge species. As described above, *E*. sp.2 and *H. stellifera* microbiomes were primarily composed of the *Nitrosopumilus*-like OTU1 and the *Nitrosopumilus* OTU3, respectively, and *Gammaproteobacteria* OTUs 2 and 7, respectively. All *Endectyon* species, i.e. *Endectyon delaubenfelsi, Endectyon* sp.1 and *Endectyon* sp.2, had the *Nitrosopumilus* OTU1 and the *Gammaproteobacteria* OTU5 as key symbionts (Fig. 7).

**Supplementary Fig. S1.**
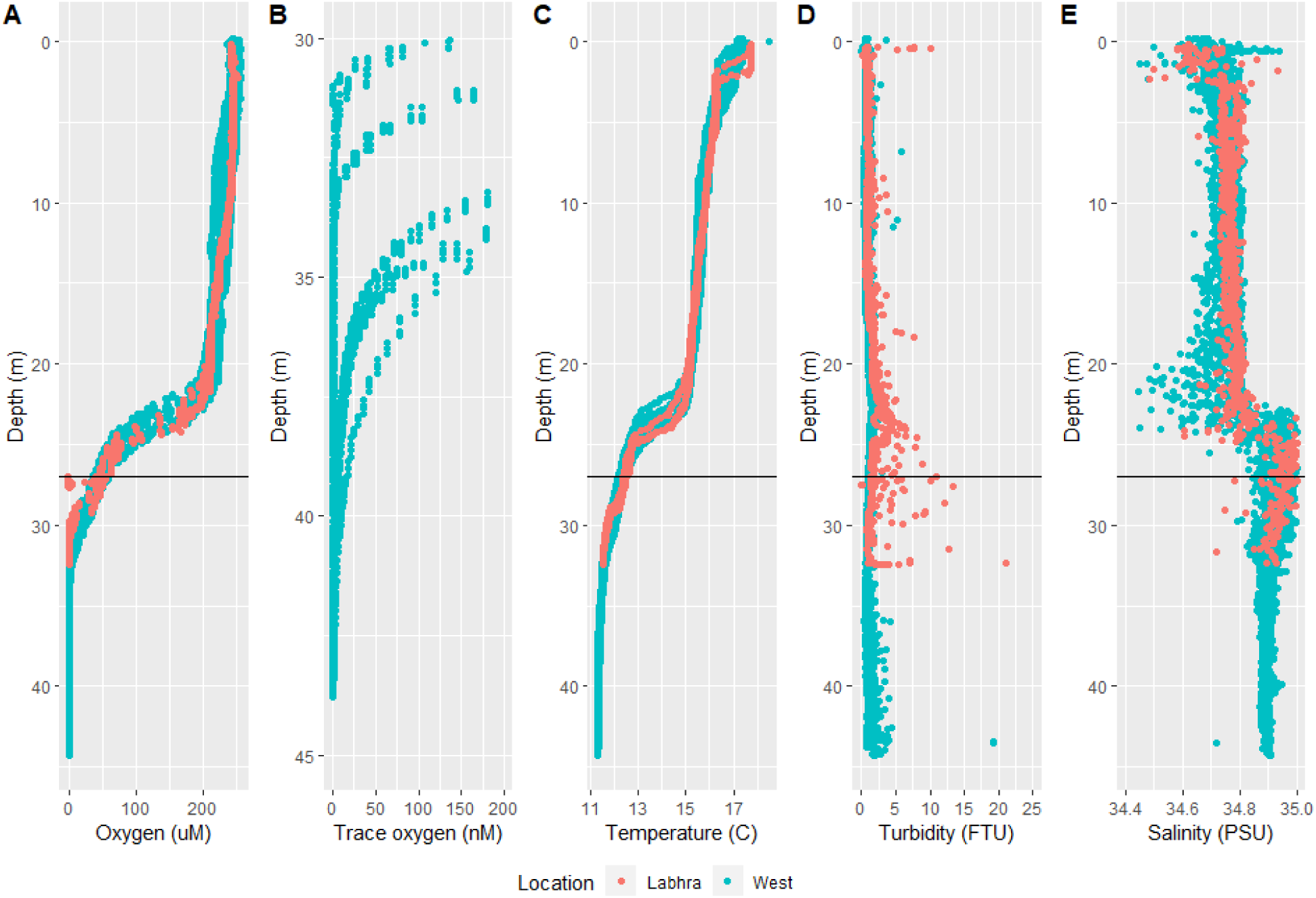
Conductivity temperature and depth (CTD) measurements during the ‘hypoxic’ conditions in July of 2019 at two sites: Labhra Cliff (red) and the Western Trough (blue). Horizontal lines present at 27 m indicate where sponges in the hypoxic condition were sampled. A. Oxygen concentrations versus depth with using the CTD’s full range (200 nM to 200 µM) oxygen probe. B. Oxygen concentrations below 200 nM measured using the trace (0 to 200 nM) oxygen probe, which registered at depths below 30 m. C. Temperature versus depth. D. Turbidity in field turbidity units (FTU). E. Salinity in practical salinity units (PSU) versus depth.

**Supplementary Fig. S2.**
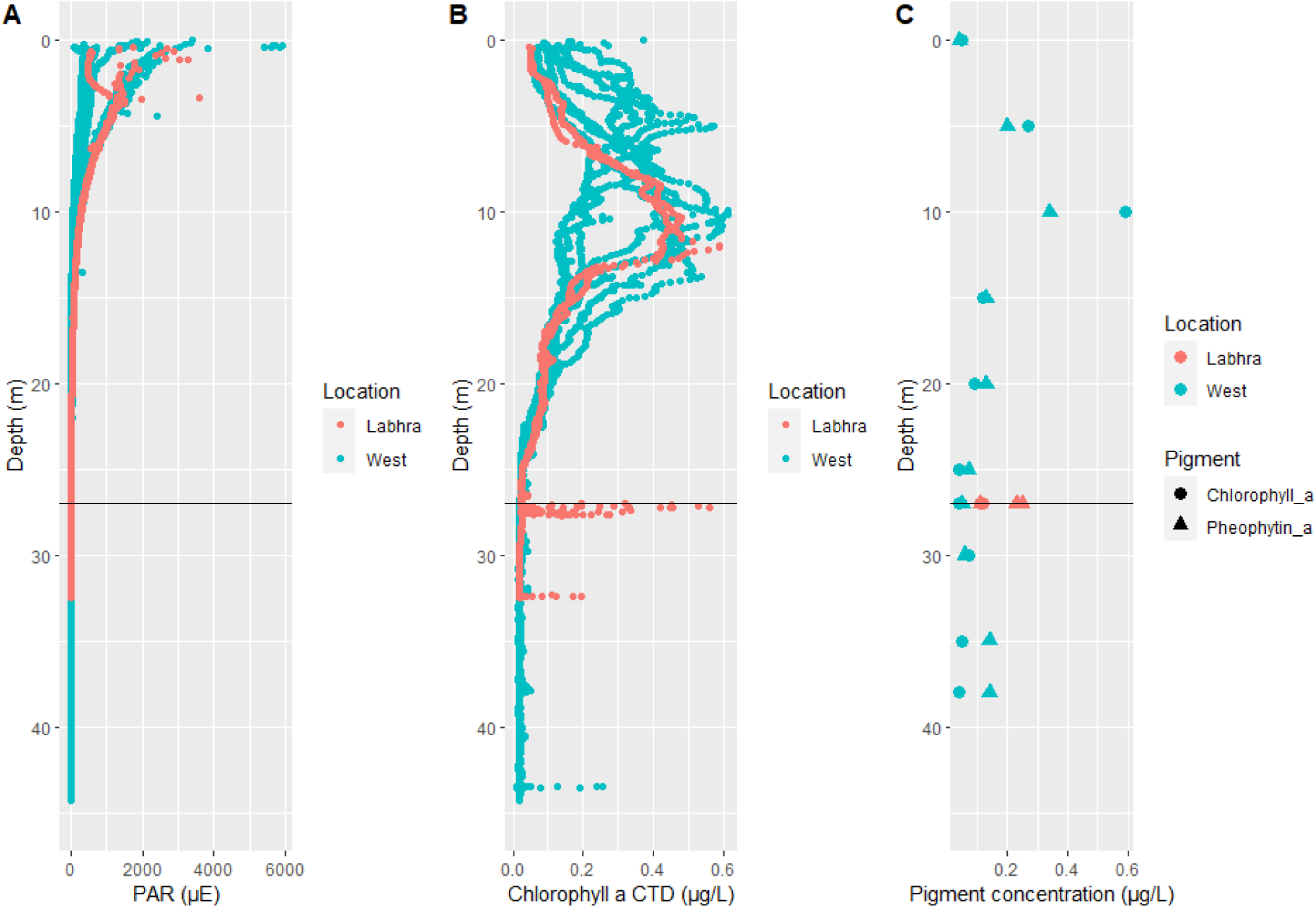
Light and pigment data from the CTD (A,B) and water samples (C) taken in July 2019. Measurements from Labhra and the western trough are listed shown in red and blue, respectively. In panel C, the concentrations of two pigments, chlorophyll a and pheophytin a are shown as circles and triangles, respectively.

**Supplementary Fig. S3.**
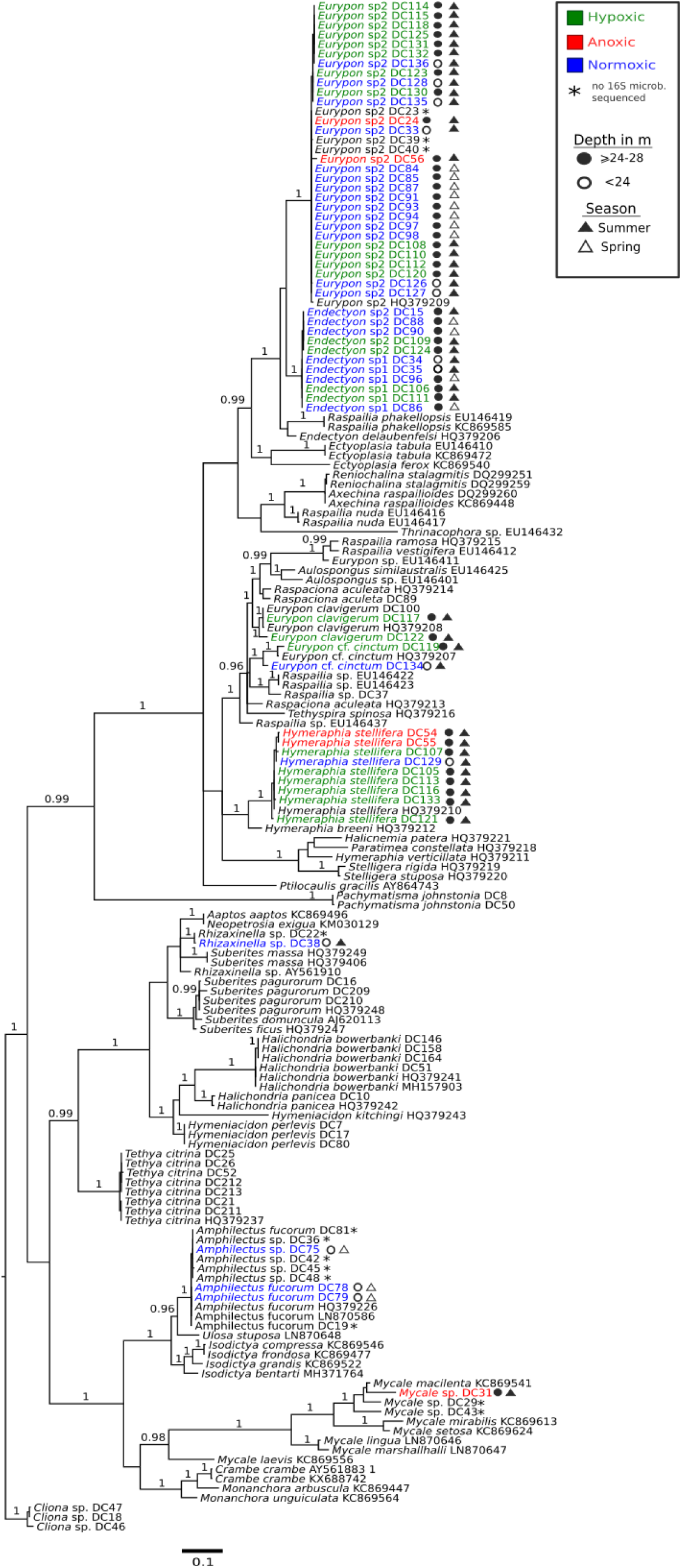
28S Bayesian Inference phylogeny of sponges from Lough Hyne (indicated by DC numbers after taxa names). Posterior Probability values are given above branches for >0.95

**Supplementary Fig. S4.**
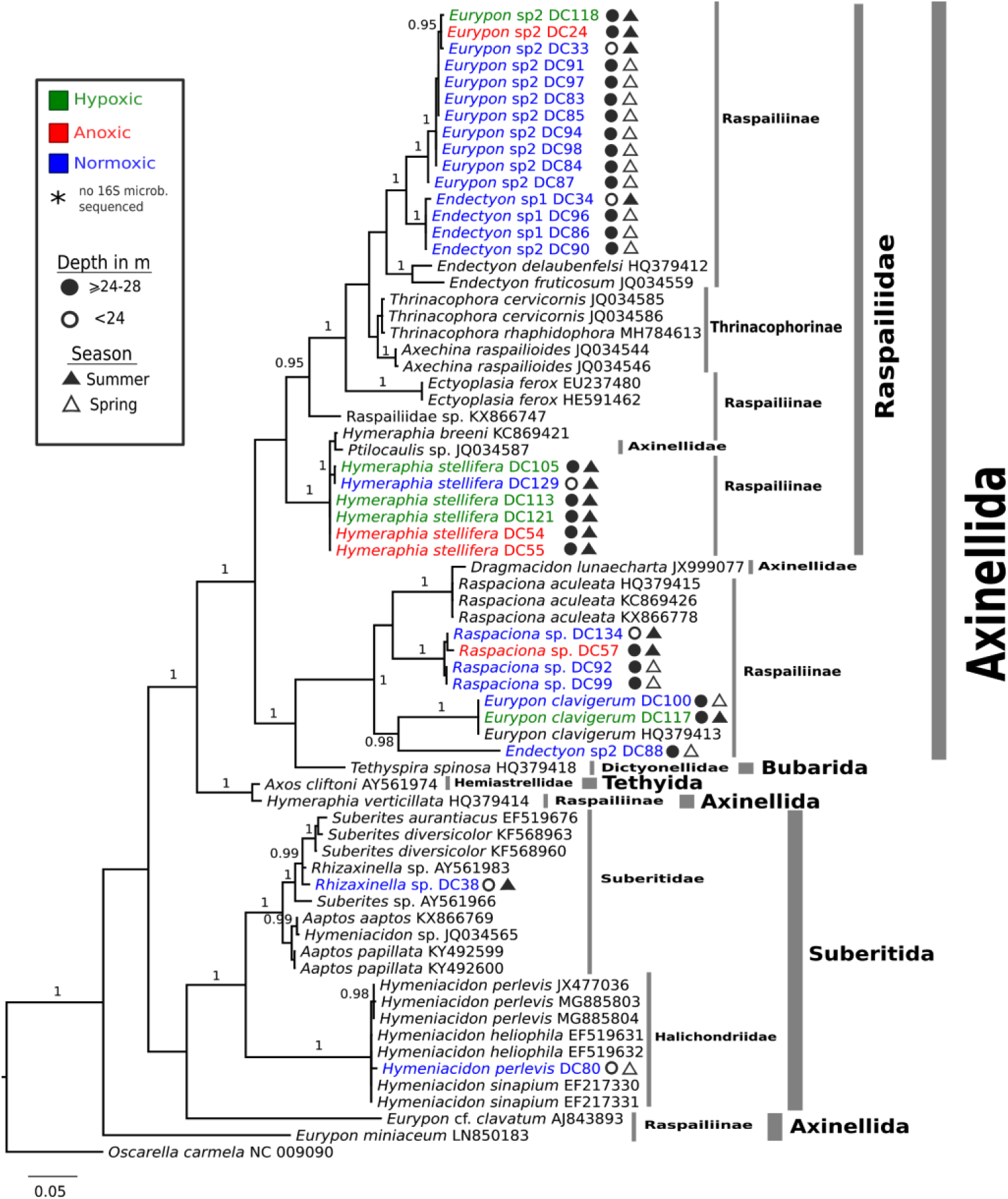
*Cox1* Bayesian Inference phylogeny of sponges from Lough Hyne (indicated by DC numbers after taxa names). Posterior Probability values are given above branches for >0.95.

**Supplementary Fig S5.**
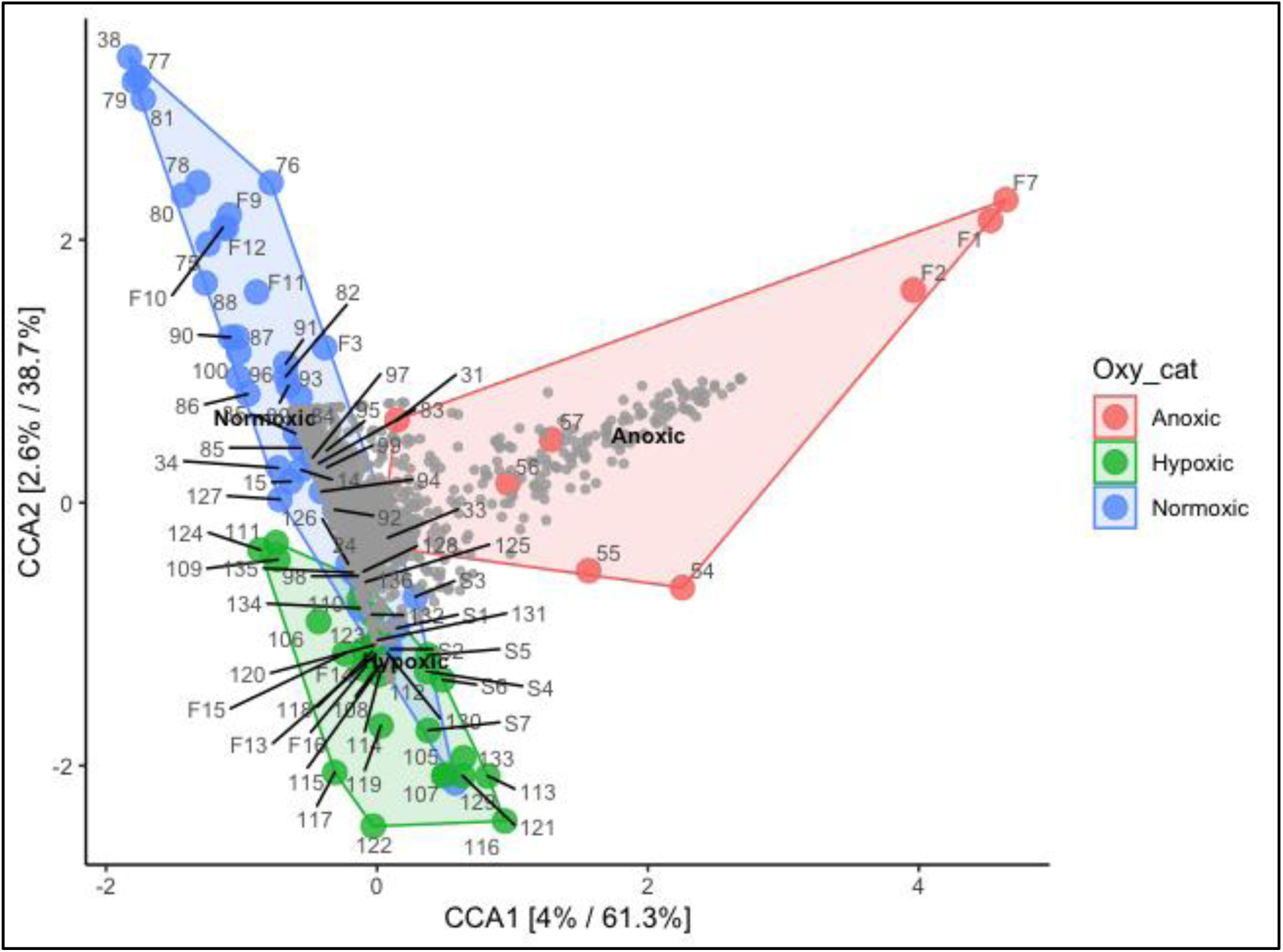
Canonical Correspondence Analysis (CCA) of all Labhra Cliff samples constrained based on oxygen condition (F=Filter, S=Sediment, ‘DC’ numbers correspond to sponge samples).

**Supplementary Table S1.**
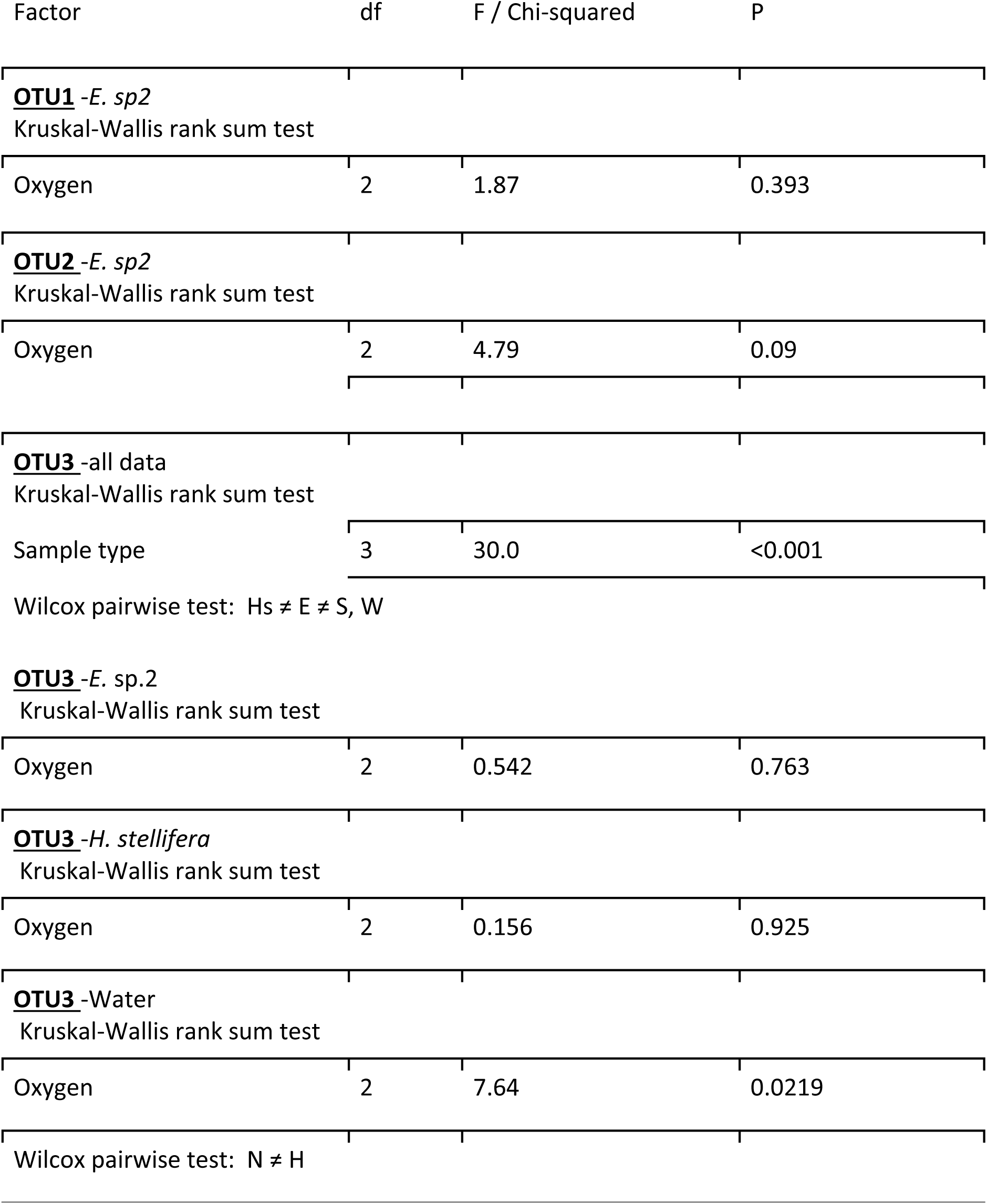
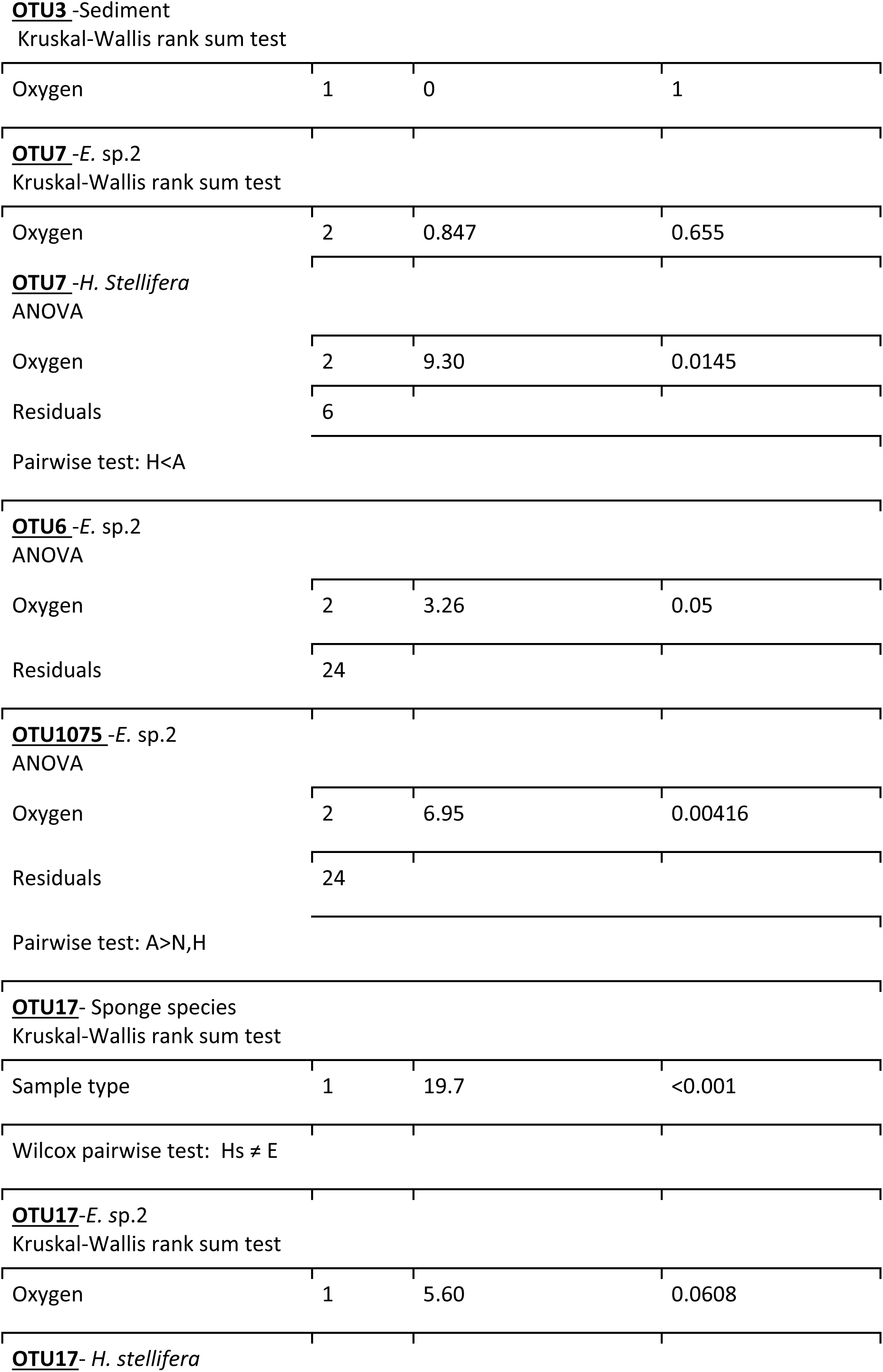
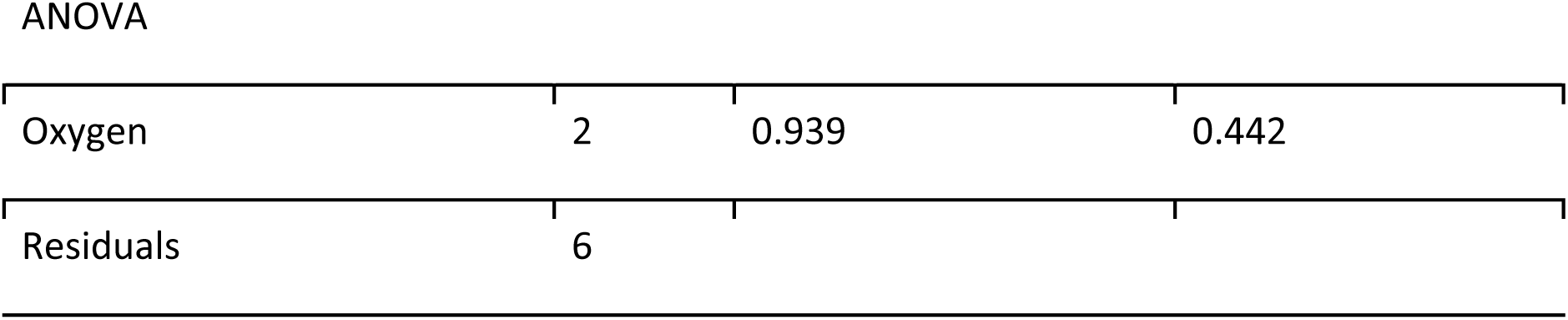
Summaries of statistical comparisons of the relative abundances of the seven most abundant OTUS within the focused subset. The type of test used is specified, and F and Chi Squared statistics are reported for parametric and nonparametric tests, respectively. Post Hoc differences are summarised using the following codes: anoxic (A), Hypoxic (H), Normoxic (N), *Eurypon* Sp2 (E), *Hymeraphia stellifera* (Hs), water (W), and sediment (S).

| Factor | df | F / Chi-squared | P |
| --- | --- | --- | --- |
| **<u>OTU1</u>** -*E. sp2*<br>Kruskal-Wallis rank sum test |  |  |  |
| Oxygen | 2 | 1.87 | 0.393 |
| **<u>OTU2</u>** -*E. sp2*<br>Kruskal-Wallis rank sum test |  |  |  |
| Oxygen | 2 | 4.79 | 0.09 |
| **<u>OTU3</u>** -all data<br>Kruskal-Wallis rank sum test |  |  |  |
| Sample type | 3 | 30.0 | <0.001 |
| Wilcox pairwise test: Hs ≠ E ≠ S, W | | | |
| **<u>OTU3</u>** -*E. sp.2*<br>Kruskal-Wallis rank sum test |  |  |  |
| Oxygen | 2 | 0.542 | 0.763 |
| **<u>OTU3</u>** -*H. stellifera*<br>Kruskal-Wallis rank sum test |  |  |  |
| Oxygen | 2 | 0.156 | 0.925 |
| **<u>OTU3</u>** -Water<br>Kruskal-Wallis rank sum test |  |  |  |
| Oxygen | 2 | 7.64 | 0.0219 |
| Wilcox pairwise test: N ≠ H | | | |
| Oxygen | 1 | 0 | 1 |
| Oxygen | 2 | 0.847 | 0.655 |
| Oxygen | 2 | 9.30 | 0.0145 |
| Residuals | 6 |  |  |
| Oxygen | 2 | 3.26 | 0.05 |
| Residuals | 24 |  |  |
| Oxygen | 2 | 6.95 | 0.00416 |
| Residuals | 24 |  |  |
| Sample type | 1 | 19.7 | <0.001 |
| Oxygen | 1 | 5.60 | 0.0608 |
| ANOVA |  |  |  |
| Oxygen | 2 | 0.939 | 0.442 |
| Residuals | 6 |  |  |

**Supplementary Fig. S6.**
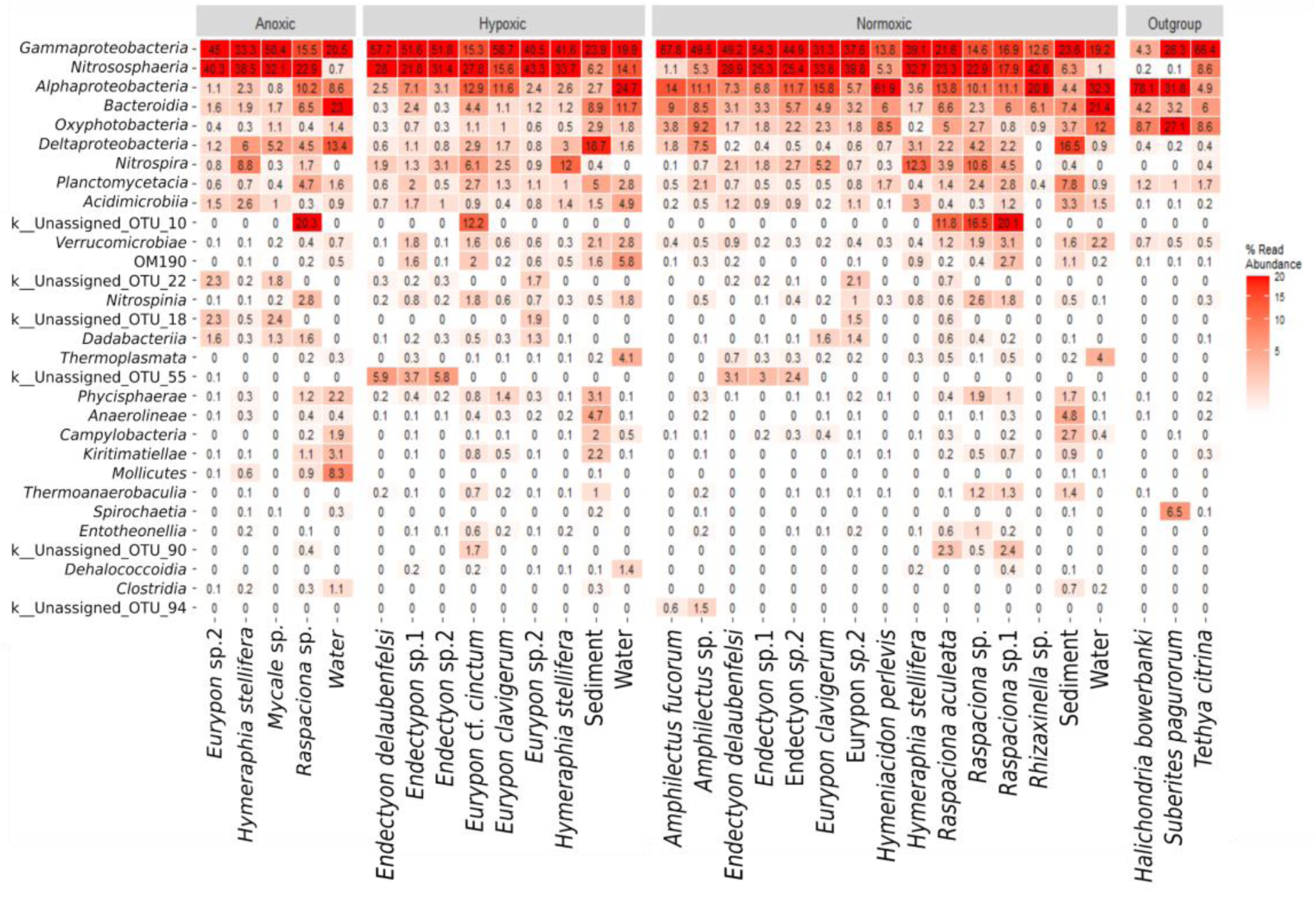
Heatmap of relative abundances of microbes at class level taxonomy (Top 30 most abundant classes) for all samples.

